# Hepatoblastoma exhibits a predominantly myeloid immune landscape and reveals opportunities for macrophage targeted immunotherapy

**DOI:** 10.1101/2023.06.28.546852

**Authors:** Daniëlle Krijgsman, Lianne Kraaier, Meggy Verdonschot, Stephanie Schubert, Jeanette Leusen, Evelien Duiker, Ruben de Kleine, Vincent de Meijer, Ronald de Krijger, József Zsiros, Weng Chuan Peng, Yvonne Vercoulen

**Affiliations:** Center for Molecular Medicine, University Medical Center Utrecht, Utrecht University, 3584, CX, Utrecht, the Netherlands; Center for Translational Immunology, University Medical Center, Utrecht, the Netherlands; Princess Máxima Center for Pediatric Oncology, Heidelberglaan 25, 3584 CS, Utrecht, the Netherlands; Department of Pathology and Medical Biology, University of Groningen, University Medical Center Groningen, Groningen, the Netherlands; Department of Surgery, Section of Hepatobiliary Surgery and Liver Transplantation, University of Groningen, University Medical Center Groningen, Groningen, the Netherlands; UCyTOF, Center for Molecular Medicine, University Medical Center Utrecht, Utrecht University, 3584, CX, Utrecht, the Netherlands

**Keywords:** Hepatoblastoma, IMC, single-cell RNA sequencing, macrophages, immune landscape, immunotherapy

## Abstract

**Background & Aims:** Hepatoblastoma (HB) is a rare form of pediatric liver cancer which is currently treated with chemotherapy and surgery. The side effects of chemotherapy pose a major problem in HB and underline the need for an alternative treatment option. We aimed to characterize the immune landscape of HB to improve our understanding of the immunologic contribution to this disease and explore immunotherapeutic options.

**Methods:** An imaging mass cytometry panel of 36 antibodies was used on tissue of treatment-naive HB (n=5), and chemotherapy-treated HB (n=3), with paired distal normal liver tissue. Immunofluorescence was used to stain HB and normal liver tissue for Kupffer cell marker MARCO. A public single-cell RNA-sequencing (scRNA-seq) dataset was analyzed consisting of 9 chemotherapy-treated HB and paired normal liver tissue.

**Results:** HB showed a heterogeneous immune landscape predominantly comprising macrophages and monocytes with high expression of immune checkpoints CD47, SIRPα, and VISTA, whereas T cells were limited. Chemotherapy increased influx of macrophages and CD8^+^ T cells in HB. Transcriptome profiling demonstrated an early activated phenotype of CD8^+^ T cells in chemotherapy-treated HB and absence of an exhaustion signature and immune checkpoint expression. Furthermore, tumor-associated macrophages had low *MARCO* expression, upregulated inflammatory markers and a high liver tissue residency score while expressing other Kupffer cell markers, such as *CD5L*, to a variable degree.

**Conclusions:** The absence of immune checkpoints and exhaustion markers in CD8^+^ T cells prohibits T cell-targeting by immune checkpoint blockade in HB patients. Instead, HB tumors contain a large myeloid compartment which provide opportunities for macrophage targeting, thereby paving the way for the development of improved treatment strategies for HB patients.

**Graphical abstract:** 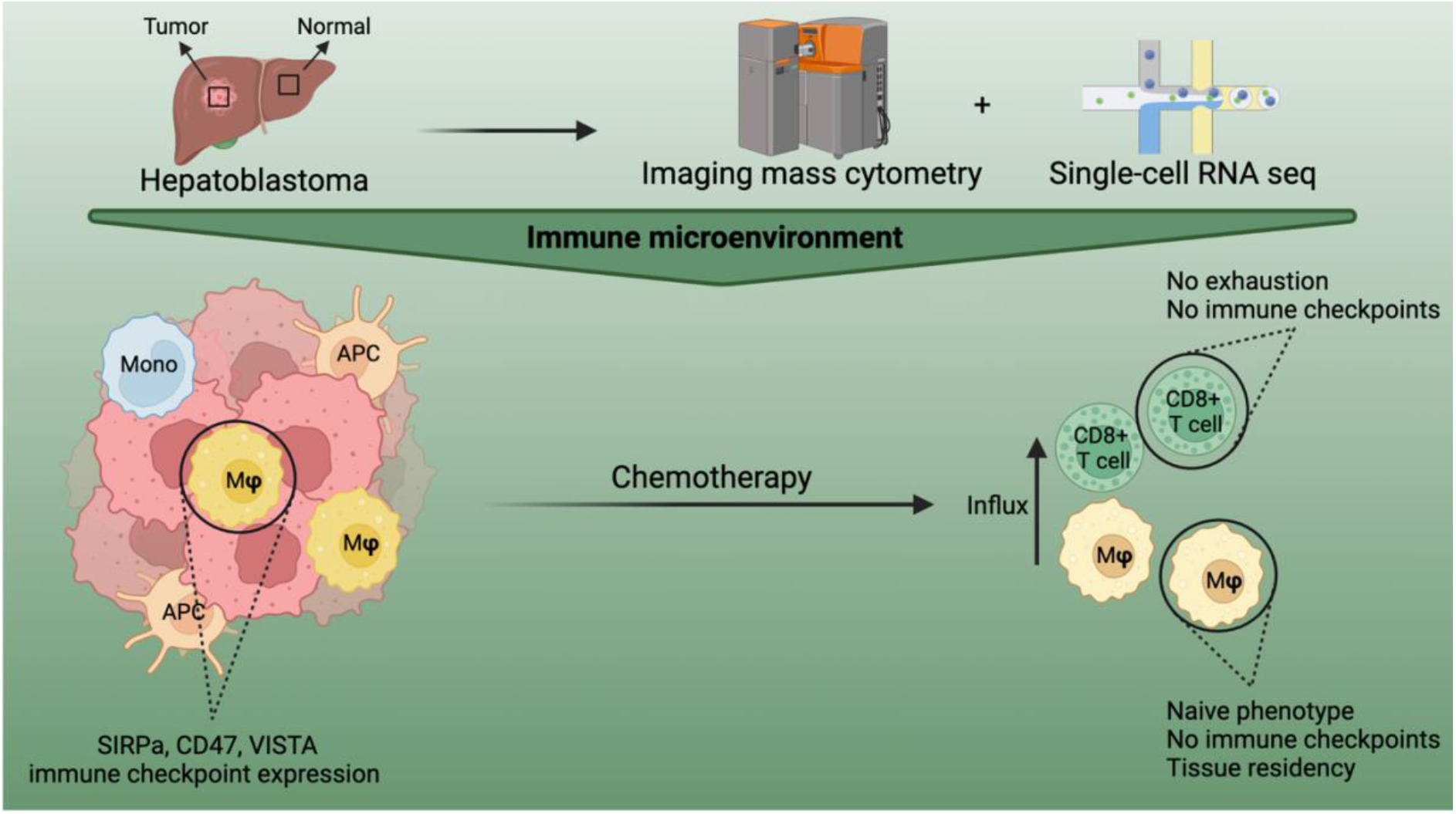

## Introduction

Hepatoblastoma (HB) is the most common primary malignant liver tumor in children, for which radical resection, in combination with chemotherapy, is the only curative treatment option ^1,2^. Although advancements in protocolised care have improved the overall survival rates for many HB patients, toxic side effects associated with chemotherapy, such as hearing loss and the development of secondary malignancies ^3,4^, can significantly impact the patient’s quality of life ^5^. Furthermore, high-risk patients with multifocal disease, or pulmonary metastases, continue to experience poor survival outcome with conventional chemotherapeutic regimens, underscoring the need for an alternative therapeutic approach ^6,7^.

One promising area of research is the development of immunotherapies, which have demonstrated success in the treatment of various malignancies ^8^. For instance, in adult hepatocellular carcinoma (HCC), exhausted T cell populations have been found to express various checkpoint molecules ^9,10^, which have led to clinical application of immune checkpoint therapy for advanced HCC. However, due to the rarity of HB, the immune landscape of this pediatric liver cancer is largely unexplored, and the feasibility of such an approach has not yet been established.

Recent studies briefly touched upon the HB immune landscape, using immunohistochemistry (IHC), bulk RNA sequencing (RNA-seq) and single-cell RNA-seq (scRNA-seq). For instance, Li et al. showed that a high density of M2-like tumor-associated macrophages (TAMs) was present in HB with embryonal differentiation using three-marker IHC and suggested that direct crosstalk between tumor cells and TAMs promotes tumor progression ^11^. Additionally, Song et al. showed, using scRNA-seq, that macrophages were enriched within chemotherapy-treated HB compared to distal normal liver tissue ^12^. These TAMs exhibited distinct transcriptional characteristics from Kupffer cells (KC), the tissue resident macrophages of the liver, and showed upregulation of several pro-tumorigenic genes. In line with these studies, Hirsch et al. observed an overall increase in macrophages upon chemotherapy treatment using bulk RNA-seq ^13^. Finally, Wang et al. showed aberrant accumulation of erythroblastic islands, consisting of VCAM1^+^ macrophages and erythroid cells, in 13 treatment-naive HB patients using scRNA-seq and five-marker immunofluorescence (IF) which inhibited dendritic cells, resulting in poor survival ^14^.

From these studies, we can conclude that macrophages seem to play an important role in HB. However, a comprehensive overview of all macrophage subsets that can be identified in HB and their role in the tumor microenvironment (TME) is currently missing. In this study, we therefore employ a synergistic approach combining single-cell spatial high-plex imaging mass cytometry (IMC) of the HB and paired distal normal liver tissue, with scRNA-seq analysis to enhance our understanding of the immune landscape organization of HB for the development of myeloid-targeting therapies.

## Materials & Methods

### Study Population and sample selection

Seven patients diagnosed with HB in the Princess Máxima Center (PMC, the Netherlands) who underwent radical surgery in the University Medical Center Groningen (UMCG, the Netherlands) were included in the present study. Treatment-naive tumor biopsies (n=4), treatment-naïve tumor resection (n=1) and chemotherapy-treated resections of tumor and paired distal normal tissue (n=3) were obtained. Extensive clinicopathological data is given in Table 1. All patient samples and clinical data were obtained after approval by the Biobank and Data Access Committee of the PMC (PMCLAB2020-107). All patients and/or their legal representatives signed informed consent. Two 500×500 µm regions of interest (ROIs) were annotated blindly for each sample by an expert pathologist (Ronald de Krijger). In normal tissues, one ROI was annotated in the liver parenchyma, and one containing the portal triad.

**Table 1:**
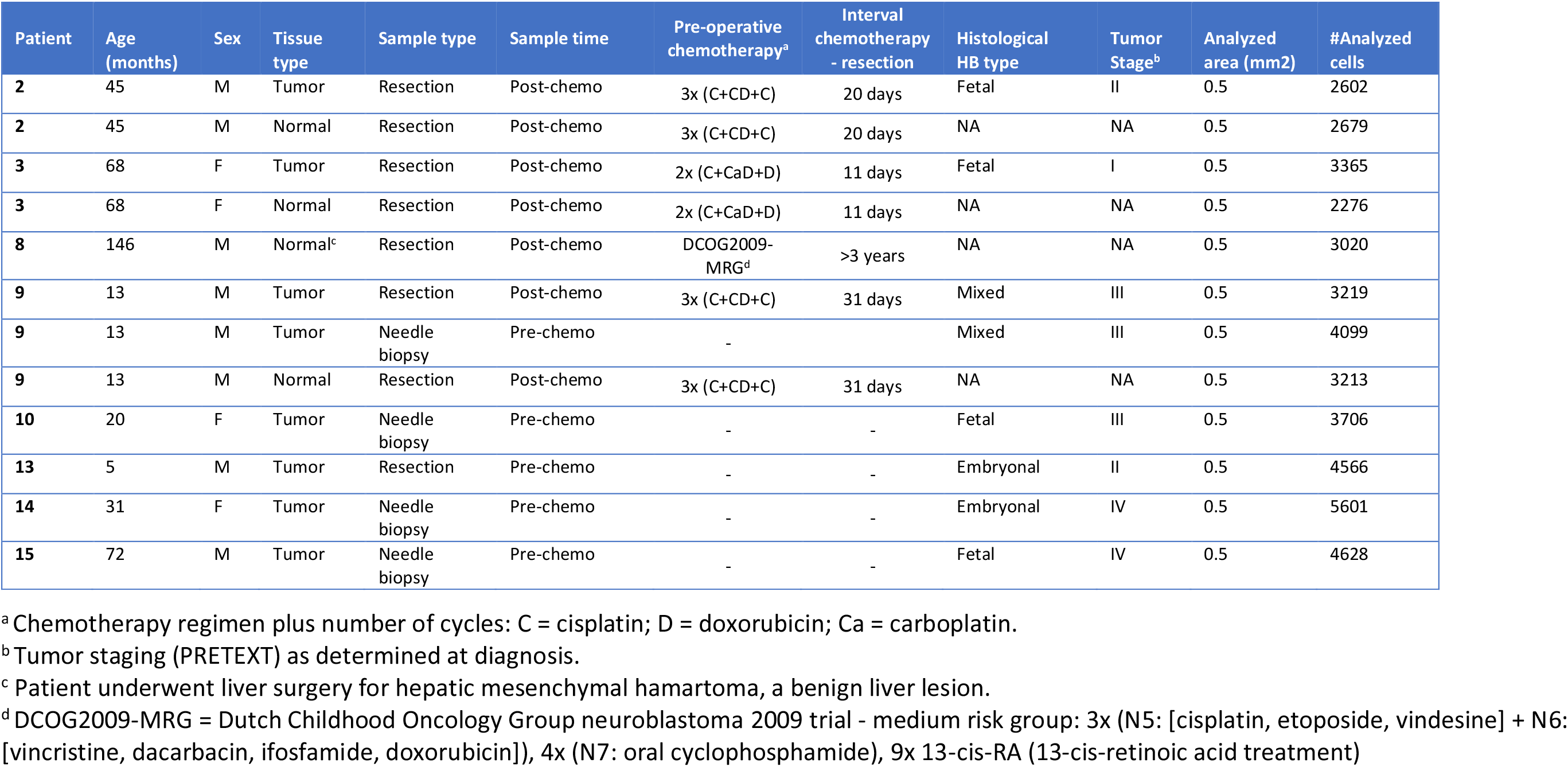
Clinical data and cell segmentation information per patient.

| Patient | Age (months) | Sex | Tissue type | Sample type | Sample time | Pre-operative chemotherapy<sup>a</sup> | Interval chemotherapy - resection | Histological HB type | Tumor Stage<sup>b</sup> | Analyzed area (mm2) | #Analyzed cells |
| --- | --- | --- | --- | --- | --- | --- | --- | --- | --- | --- | --- |
| 2 | 45 | M | Tumor | Resection | Post-chemo | 3x (C+CD+C) | 20 days | Fetal | II | 0.5 | 2602 |
| 2 | 45 | M | Normal | Resection | Post-chemo | 3x (C+CD+C) | 20 days | NA | NA | 0.5 | 2679 |
| 3 | 68 | F | Tumor | Resection | Post-chemo | 2x (C+CaD+D) | 11 days | Fetal | I | 0.5 | 3365 |
| 3 | 68 | F | Normal | Resection | Post-chemo | 2x (C+CaD+D) | 11 days | NA | NA | 0.5 | 2276 |
| 8 | 146 | M | Normal<sup>c</sup> | Resection | Post-chemo | DCOG2009-MRG<sup>d</sup> | >3 years | NA | NA | 0.5 | 3020 |
| 9 | 13 | M | Tumor | Resection | Post-chemo | 3x (C+CD+C) | 31 days | Mixed | III | 0.5 | 3219 |
| 9 | 13 | M | Tumor | Needle biopsy | Pre-chemo | - | - | Mixed | III | 0.5 | 4099 |
| 9 | 13 | M | Normal | Resection | Post-chemo | 3x (C+CD+C) | 31 days | NA | NA | 0.5 | 3213 |
| 10 | 20 | F | Tumor | Needle biopsy | Pre-chemo | - | - | Fetal | III | 0.5 | 3706 |
| 13 | 5 | M | Tumor | Resection | Pre-chemo | - | - | Embryonal | II | 0.5 | 4566 |
| 14 | 31 | F | Tumor | Needle biopsy | Pre-chemo | - | - | Embryonal | IV | 0.5 | 5601 |
| 15 | 72 | M | Tumor | Needle biopsy | Pre-chemo | - | - | Fetal | IV | 0.5 | 4628 |

<sup>a</sup> Chemotherapy regimen plus number of cycles: C = cisplatin; D = doxorubicin; Ca = carboplatin.
<sup>b</sup> Tumor staging (PRETEXT) as determined at diagnosis.
<sup>c</sup> Patient underwent liver surgery for hepatic mesenchymal hamartoma, a benign liver lesion.
<sup>d</sup> DCOG2009-MRG = Dutch Childhood Oncology Group neuroblastoma 2009 trial - medium risk group: 3x (N5: [cisplatin, etoposide, vindesine] + N6: [vincristine, dacarbacin, ifosfamide, doxorubicin]), 4x (N7: oral cyclophosphamide), 9x 13-cis-RA (13-cis-retinoic acid treatment)

### Antibody conjugation

The MaxPar labeling kit (Standard BioTools Inc.) was used to conjugate carrier-free antibodies to metal isotopes for IMC following manufacturer’s protocol. Table S1 summarizes the antibodies used in this study.

### IMC sample preparation

Four µm formalin-fixed paraffin-embedded (FFPE) tissue sections were cut and placed on glass slides (Klinipath) and baked for an hour at 60 °C. Tissue sections were then stained as previously described ^15,16^. Upon Fc-block, tissue sections were incubated overnight with anti-SIRPα in Tris-buffered saline with 0.1% Tween® 20 detergent (TBST) containing 0.5% bovine serum albumin (BSA) in a humidified chamber at 4 °C. The next day, samples were washed 3 times for 5 min in TBST and incubated with secondary rabbit-161Dy antibody in TBST containing 0.5% BSA for 1 hour at room temperature (RT).

Tissue sections were then washed 5 min for 3 times in TBST, followed by overnight incubation at 4 °C at 4 °C with a mix of metal-labeled antibodies in TBST containing 0.5% BSA as listed in Table S1. Upon washing twice with TBST, slides were stained with 500 times diluted DNA intercalator Ir193 (Standard BioTools Inc.) and 5 µg/ml DAPI (Sigma-Aldrich) for 1 hour at RT. Slides were then imaged by fluorescence microscopy, followed by IMC as previously described ^16^.

### IMC single-cell data generation and downstream analyses

The MATISSE pipeline was used to process fluorescence and IMC images to single-cell data as previously described ^15,16^. A tissue microarray (TMA) was included on every glass slide, containing tissue of different origins, which were also stained and ablated and used to inspect for inter-experimental variation between slides. 99^th^ percentile normalization was used for data normalization. Single-cell data was log1p transformed for clustering and expression analyses in R (version 4.1.2). First, all cells across samples were integrated by FastMNN mutual nearest neighbor correction (batchelor package), based on the mean expression per cell of: CD45, MPO, CD15, CD16, CD68, CD20, CD4, CD3, CD8a, CD14, HLA-DR, and CD11b. Uniform manifold approximation and projection (UMAP) was computed using the umap package. Clusters containing immune cells were selected based on CD45 expression (Figure S1). Thereafter, the immune cells from normal and tumor samples were re-clustered separately using the markers described above. B cells and neutrophils were manually gated based on CD20 and CD15 expression, respectively. Tumor-derived T cells were subclustered into CD4^+^ and CD8^+^ T cells via manual gating based on CD8 expression. CytoMAP was used for neighborhood analysis using a raster scanned approach with a radius of 50 µm ^17^. A cut-off of >0.45 was used to define cell interactions based on the Pearson correlation tests.

### Single cell RNA sequencing analyses

Single-cell transcriptome data of HB and distal normal liver was accessed through the GEO database (accession number GSE186975) ^12^. Raw UMI-collapsed read-count data was analyzed through Seurat (version 4.3.0) in R (version 4.1.2) ^18^. Contamination by ambient RNA was estimated and removed by DecontX (celda version 1.10.0) using default settings ^19^. Low-quality cells were filtered by removing cells with <500 genes or >20% mitochondrial genes. A maximum threshold for number of genes was set for individual samples. Separate workflows were implemented for dimensionality reduction and gene expression analyses. For dimensionality reduction and clustering, normalization was performed by SCTransform (version 0.3.5), and cell-cycle and related genes were identified as previously described ^20^. Different batches were integrated using fastMNN batch correction (SeuratWrappers version 0.3.1), with cell cycle and related genes removed from the integration features. Dimensionality reduction and clustering were performed using 40 PC and a resolution of 0.5, yielding 22 clusters. For gene expression analyses, counts were normalized and log-transformed by Seurat’s “LogNormalize” method. Differentially expressed genes (DEG) for each cluster were identified by Seurat’s FindAllMarkers and facilitated cell type identification.

After subsetting of immune cell types, dimensionality reduction and clustering were recomputed on the fastMNN-corrected data. Macrophages and monocytes clusters were further subsetted and processed as described above. DEGs were defined using cutoffs of Log2FC > 1 and p-value <0.05. The tissue resident macrophage signatures were generated as previously described ^21^. Briefly, the top 25 DEGs of TLF^+^ macrophages were taken from each organ and converted to human gene orthologs using g:Profiler Orthology search. Gene signature scores were generated using Seurat’s AddModuleScore.

### Immunofluorescent staining of Kupffer cells

Four µm FFPE tissue sections were cut and placed on glass slides (SuperFrost® Plus). Tissue sections were deparaffinized and rehydrated and heat-induced antigen retrieval was performed in citrate buffer (10mM sodium citrate dihydrate, pH 6.0) for 20 min. Samples were encircled using a PAP pen (Sigma-Aldrich) and incubated with blocking buffer (5% normal donkey serum [Jackson ImmunoResearch] in 0.1% Triton X-100 in PBS [PBS-Tx]) for 1 hour at RT. Thereafter, tissue sections were incubated with primary antibodies (Table S2) diluted in blocking buffer in a humidified chamber overnight at 4 °C. The next day, samples were washed 3 times with 0.1% PBS-Tx and incubated with secondary antibodies (Table S2) diluted in PBS and for 1 hour at RT. Tissue sections were then washed 3 times in PBS, followed by incubation with DAPI at 5 µg/ml for 5 min at RT. Finally, sections were washed 3 more times and mounted in 80% glycerol in PBS and a #1.5 coverslip. Images were acquired with a 40X water immersion objective on a Leica DMi8 Thunder widefield microscope equipped with 4 LED light sources (DAPI [395/25], FITC [475/28],Cy3 [555/28] and Cy5 [635/22]).

### Statistical analyses

Nonparametric tests were used to compare cell clusters and marker expression in treatment-naive and chemotherapy-treated samples. The Pearson correlation tests were used for neighborhood analysis. Figures were made using BioRender, FIJI (v2.9.0/1.53t) and Adobe Illustrator (v25.0).

## Results

### HB shows decreased expression of T and B cell markers compared to distal normal liver

To comprehensively characterize the TME of HB, we used a customized IMC panel of 36 antibodies on treatment-naive (5 HB), and chemotherapy-treated (3 HB, and 4 paired distal normal liver) tissue (Figure 1A,B and Table 1). Processing of images into single-cell expression data yielded a total of 42,974 cells. Principal component analysis (PCA) of median protein expression levels per ROI revealed separation between normal liver and HB samples (Figure 1C). HB showed significantly reduced expression of T cell (CD3, CD4, CD8a, FOXP3) and B cell (CD20) markers compared to distal normal liver, suggesting the lack of lymphocyte infiltration in the tumor, while the expression of myeloid (CD11b, CD33, CD68, CD163, CD16) and granulocyte (CD15, MPO) lineage markers remained similar (Figure 1D,E and Figure S2). In addition, the monocyte and sinusoidal endothelial cell marker CD14 was significantly decreased in HB.

**Figure 1:**
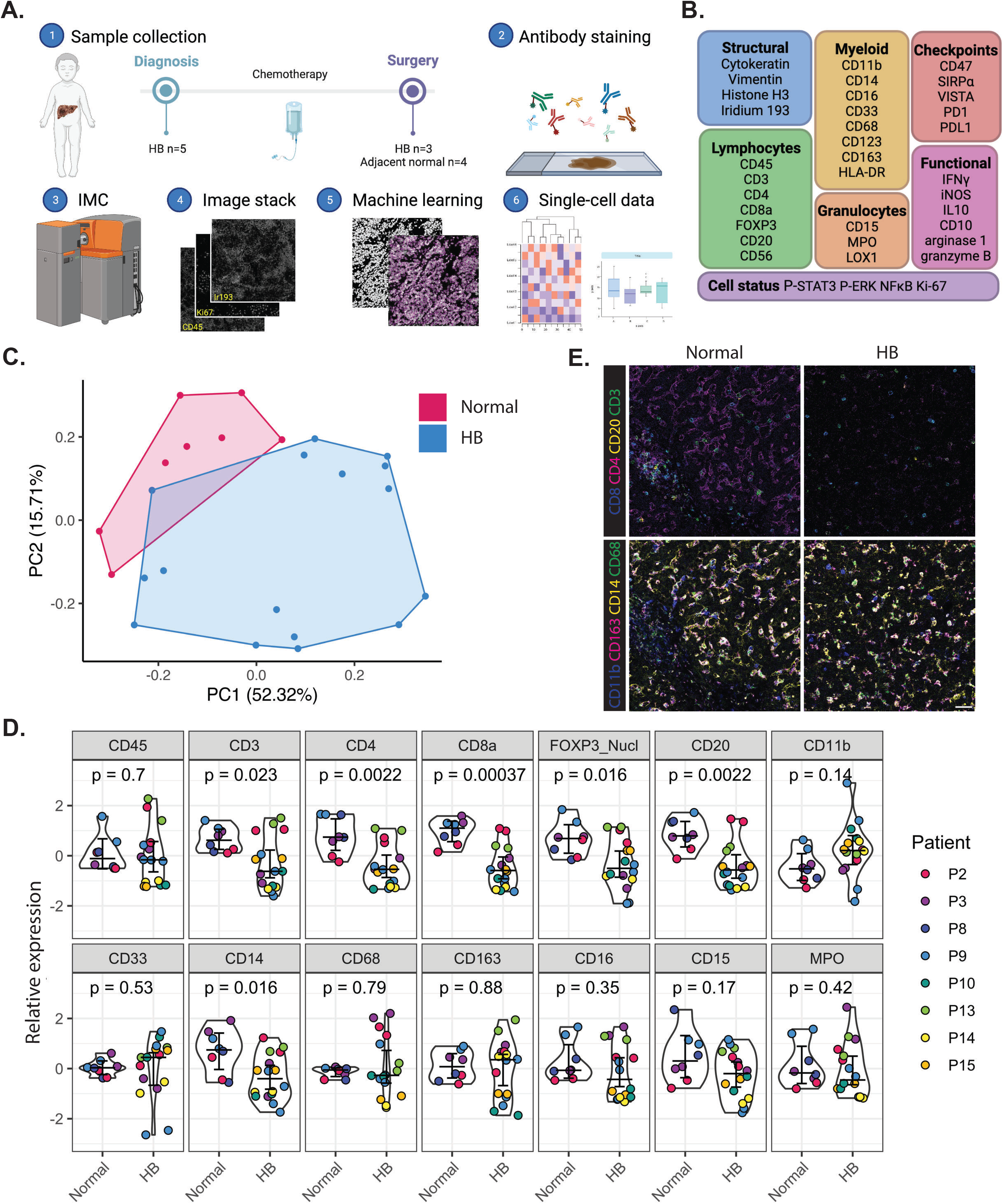
Decrease in expression of lymphocyte markers in hepatoblastoma tissue compared to distal normal liver. **A.** Illustration of the sample selection and data acquisition workflow used for CyTOF IMC. **B.** Overview of markers included in the IMC panel. **C.** PCA using the median intensity of each marker per image region as input variables. Each data point is a ROI, color-coded by sample type. **D.** Violin plots with median intensity of lineage and functional markers per patient, split for normal and HB tissue. Bars indicate the median with 95% confidence interval. **E.** Representative IMC images for HB and distal normal tissues. Scale bar is 50 μm.

### Compartmentalized immune landscape in the normal liver portal triad and parenchyma

To assess immune cell distribution in normal liver, we performed unsupervised clustering. We observed distinct clusters corresponding to immune, epithelial and stromal cells, and annotated the immune cell types based on lineage marker expression (Figure 2A,B). The mean abundance of immune cells was 35.3% (range 28.7-46.1%), which was homogeneous between patients, overall containing mostly macrophages (Figure 2C). Interestingly, cell types present in the portal triad clustered separately from parenchymal cells. Spatial evaluation revealed that ‘Antigen presenting cells (APCs)’, and ‘Macrophages & T cells’ predominantly resided in the liver portal triad compartment, while the parenchyma mainly contained ‘Macrophages’ and ‘Epithelial & stromal cells’ (Figure 2D, Figure S3). IF staining revealed that nearly all macrophages in the liver parenchyma were CD68^+^MARCO^+^ KCs, while the portal triad contained CD68^+^MARCO^-^ macrophages (Figure 2E). Unbiased neighborhood analysis also showed significant compartmentalization of ‘Macrophages (KCs)’, and ‘Macrophages and T cells’ (Figure 2F). The ‘Macrophages (KCs)’ cluster in the parenchyma showed a suppressive phenotype characterized by high CD163, CD16, iNOS and arginase-1 expression (Figure S4A,C), whereas the ‘Macrophages & T cell’ cluster in the portal triad showed high expression of HLA-DR, indicating a role in antigen presentation. Both macrophage clusters expressed the myeloid immune checkpoints SIRPα, CD47 and VISTA. Furthermore, T cells showed moderate granzyme B, IFNγ, and Ki-67 expression, indicating an activated phenotype, without expression of immune checkpoints PD1 and PDL1 (Figure S4B,D). Together, the immune landscape in normal liver is highly structured, with clustering of APCs and macrophages in the portal triad, and abundant KC and other immune subsets dispersed throughout the parenchyma.

**Figure 2:**
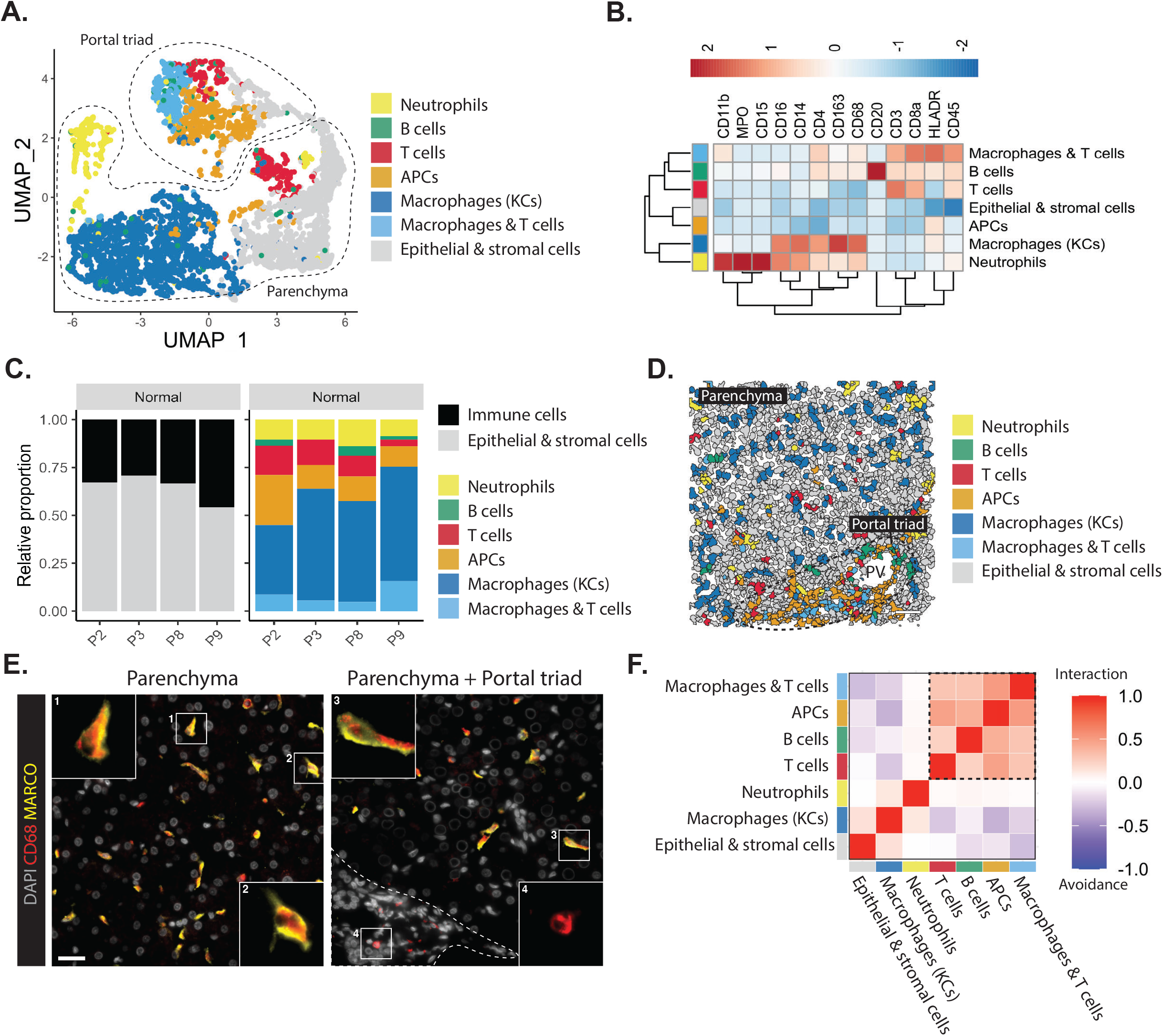
Normal liver tissue is predominated by myeloid cell types and shows a compartmentalized immune landscape. **A.** UMAP of single cells of all ROIs from normal liver tissue using lineage marker intensities, color-coded by annotated cell lineages. **B.** Hierarchical clustering heatmap of the median marker intensity per annotated cell type. **C.** Stacked bar plot representing the proportions of immune cells per patient (left), and the proportion of immune cell subsets (right). **D.** Representative IMC images. **E.** Single-cell segmented cells, annotated with cluster names. Scale bar is 50 μm. Dotted line indicates portal triad. PV: portal vein. **F.** Immunofluorescent staining of normal liver tissue. Scale bar is 25 μm. Dotted line indicates portal triad. **G.** Heatmap showing the correlation of spatial localization of cell types in normal liver tissue. Pearson correlation tests indicate avoidance (blue) or interaction (red) of cell types.

### Identification of naive and suppressive myeloid subsets in HB

HB samples displayed a heterogeneous immune cell infiltrate (Figure 3A,B), characterized by the identification of more diverse cell subsets compared to normal samples. The mean abundance of immune cells varied between patients (∼10-50%) but was significantly (p=0.011) elevated in chemotherapy-treated (47.5%) compared to treatment-naive (22.7%) HB (Figure 3C). All tumor tissues showed a predominantly myeloid infiltrate with a high abundance of either ‘APCs’, monocyte or macrophage clusters. We identified two macrophage subsets: ‘M2-like macrophages’ and ‘Infiltrating macrophages’. ‘M2-like macrophages’ displayed high levels of suppressive markers iNOS, arginase-1 and MPO, low expression of typical M2 marker CD163, but expressed myeloid immune checkpoints SIRPα, CD47 and VISTA (Figure S4). ‘Infiltrating macrophages’ showed mixed expression of M1-(iNOS) and M2-related (arginase-1, IL-10, CD163) markers, and lacked immune checkpoints, suggesting a naive phenotype as observed in infiltrating macrophages that have not yet been exposed to any pro- or anti-inflammatory stimuli ^22^. Furthermore, two monocyte subsets were identified: ‘Classical (CL) monocytes’ and ‘Non-classical (NC) monocytes & T cells’. ‘CL monocytes’, also called inflammatory monocytes, displayed low expression of suppressive markers iNOS, arginase-1 and IL-10, and lacked immune checkpoints (Figure S4). By contrast, ‘NC monocytes & T cells’ highly expressed the immune checkpoints SIRPα, CD47 and VISTA, and suppressive markers arginase-1 and IL-10, but not iNOS (Figure S4).

**Figure 3:**
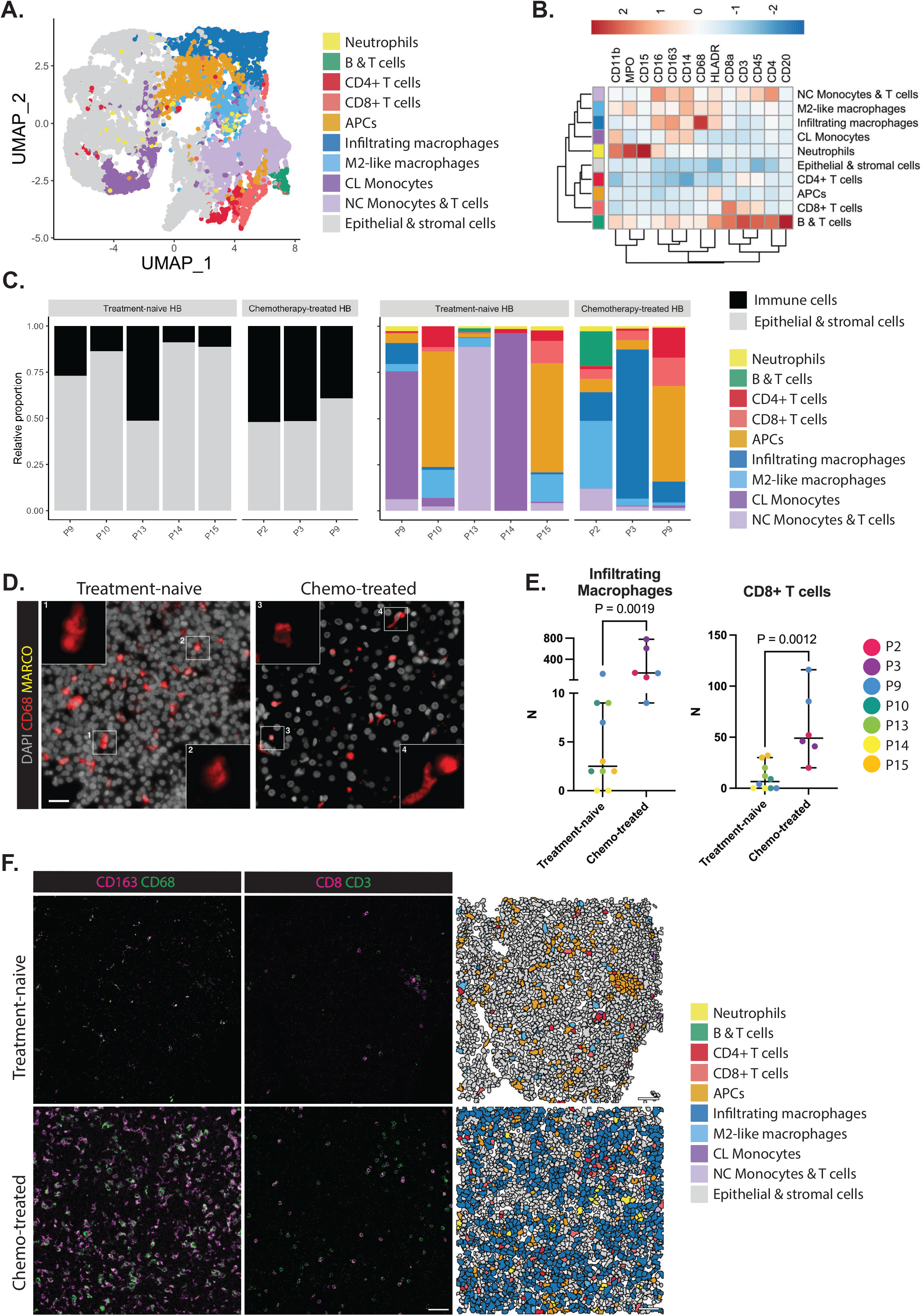
Chemotherapy induces macrophage and CD8^+^ T cell influx in hepatoblastoma. **A.** UMAP of single cells of treatment-naive and chemotherapy-treated HB tissues using lineage marker intensities, color-coded by annotated cell lineages. **B.** Hierarchical clustering heatmap of the median marker intensity per annotated cell type. Markers used for dimensionality reduction are shown. **C.** Stacked bar plot representing the proportions of immune cells per patient (left), and the proportion of immune cell subsets (right). **D.** Immunofluorescent staining of treatment-naive (left) and chemotherapy-treated (right) HB tissues. Scale bar is 25 μm. **E.** Scatter plots of the number of cells per annotated cell cluster, stratified by treatment status. Each dot is an ROI, color-coded for patient ID. The bars indicate the median with 95% confidence interval. **F.** Representative IMC images of treatment-naive and chemotherapy-treated HB tissues (left), with single-cell segmented images (right) annotated with the names of specific cell clusters in colors. Scale bar is 50 μm.

IF staining revealed that most macrophages in HB have a CD68^+^MARCO^-^ phenotype (Figure 3D). Although suppressive iNOS^+^arginase-1^+^ macrophages are known for depletion of essential amino acids required for T cell proliferation and activation ^23,24^, our IMC analysis indicated an early inflammatory phenotype of CD8^+^ T cells in HB (Figure S5), characterized by moderate granzyme B and Ki-67 expression, but absence of phosphorylated ERK (P-ERK). CD4^+^ and CD8^+^ T cells in HB lacked immune checkpoints (Figure S5). Together, we identified both naive and suppressive macrophage subsets in HB, and yet cytotoxic T cells in the TME displayed an early activated phenotype.

### Chemotherapy increases naive macrophages and CD8^+^ T cells in HB

Next, we evaluated the effect of chemotherapy on the HB immune cell subset infiltrate. As expected, the fraction of immune cells increased after chemotherapy whereas the fraction of epithelial and stromal cells decreased (Figure S6). ‘Infiltrating macrophages’ were significantly increased in HB after chemotherapy (Figure 3E,F), whereas ‘M2-like macrophages’ were unaffected (Figure S6,S7). While numbers of CD8^+^ T cells increased after chemotherapy (Figure 3E), the CD4^+^ T cell population remained stable (Figure S6). No differences in abundance were observed regarding other cell types such as neutrophils, monocytes, and APCs (Figure S6).

### Chemotherapy disrupts immune landscape tissue organization in HB

Next, we performed a neighborhood analysis to investigate spatial cell interactions in treatment-naive HB and chemotherapy-treated HB samples. First, the ‘NC Monocytes & T cells’ and ‘M2-like macrophages’ co-localized in treatment-naive HB (Figure 4A,B), indicating interaction between suppressive myeloid subsets. Furthermore, co-localization was observed between lymphocyte subsets including CD4^+^ T cells, CD8^+^ T cells, and APCs (Figure 4A,B), suggesting T cell activation via antigen presentation by APCs. By contrast, we did not observe the same co-localization in chemotherapy-treated HB (Figure 4C), likely due to disruption of the immune landscape and tissue organization following treatment. The only interaction identified in chemotherapy-treated HB, between ‘NC Monocytes & T cells’, and ‘B & T cells’, was restricted to one HB sample containing a tertiary lymphoid structure surrounded by monocytes (Figure 4C,D).

**Figure 4:**
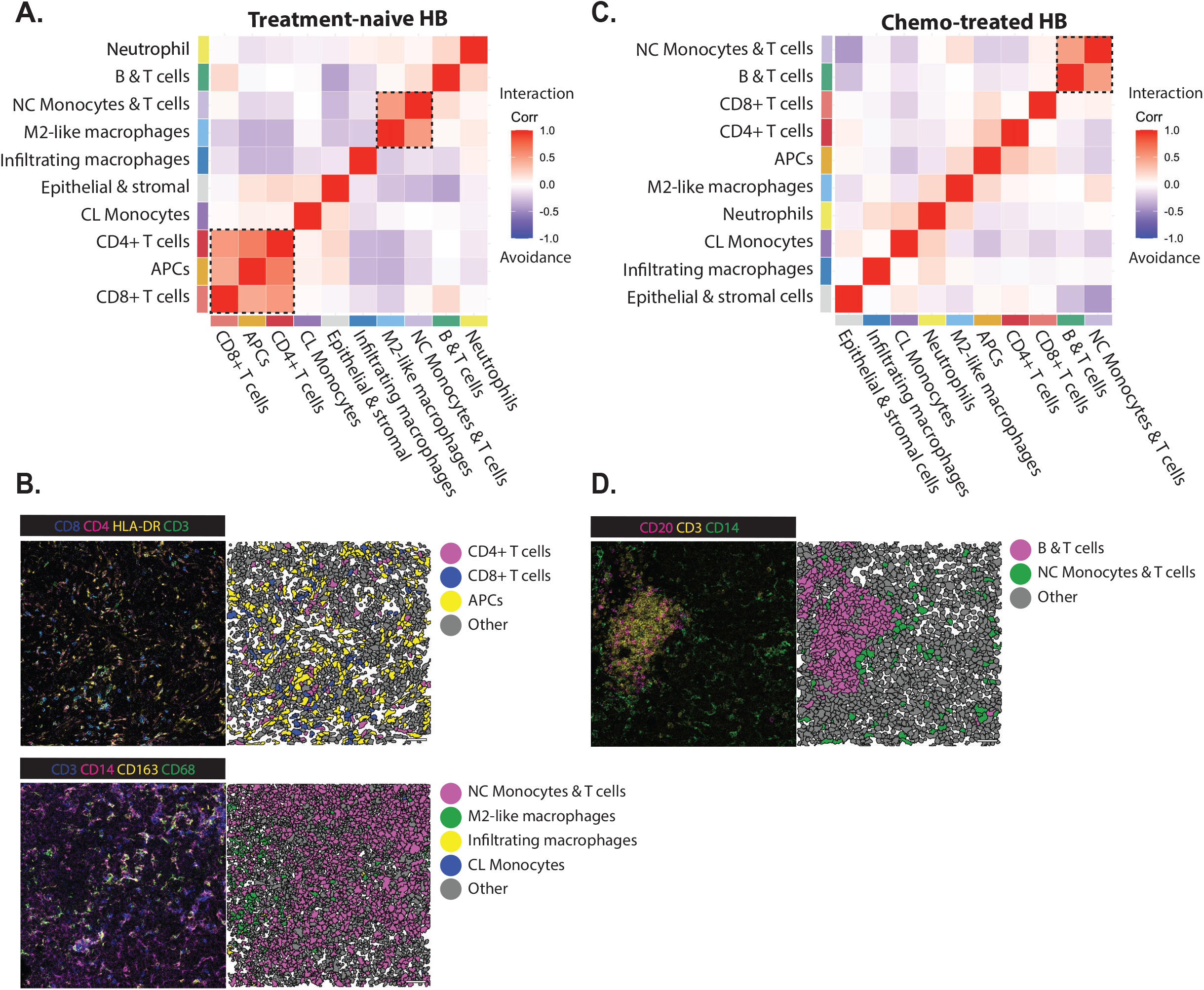
Neighborhood analysis of hepatoblastoma tissues reveals disrupted APC-T cell interactions after chemotherapy. Correlation heatmap showing the correlation of spatial localization of cell types within 50 μm raster-based defined neighborhoods in treatment-naive (**A.**) and chemotherapy-treated HB tissues (**B.**). Pearson correlation tests indicate avoidance (blue) or interaction (interaction) of cellular localization. **C.** Representative IMC images of treatment-naive HB tissues (left) with single-cell segmented cells of the same tissue (right) annotated with the names of specific cell clusters in colors. **D.** Representative IMC images of chemotherapy-treated HB tissues (left) with single-cell segmented cells of the same tissue (right) annotated with the names of specific cell clusters in colors. Scale bars are 50 μm.

### Transcriptional profiling demonstrates presence of naive and early activated T cells in HB

To further understand the immune populations identified by IMC, we analyzed recently published scRNA-seq data, consisting of 9 chemotherapy-treated HB and distal normal tissue ^12^. We focused on the immune cell types that were analyzed in our IMC data (Figure 5A-C and Figure S8A,B). Consistent with IMC, the relative distribution of immune populations per patient demonstrated a predominantly myeloid landscape and few lymphocytes in normal liver and HB tissues (Figure 5B).

**Figure 5:**
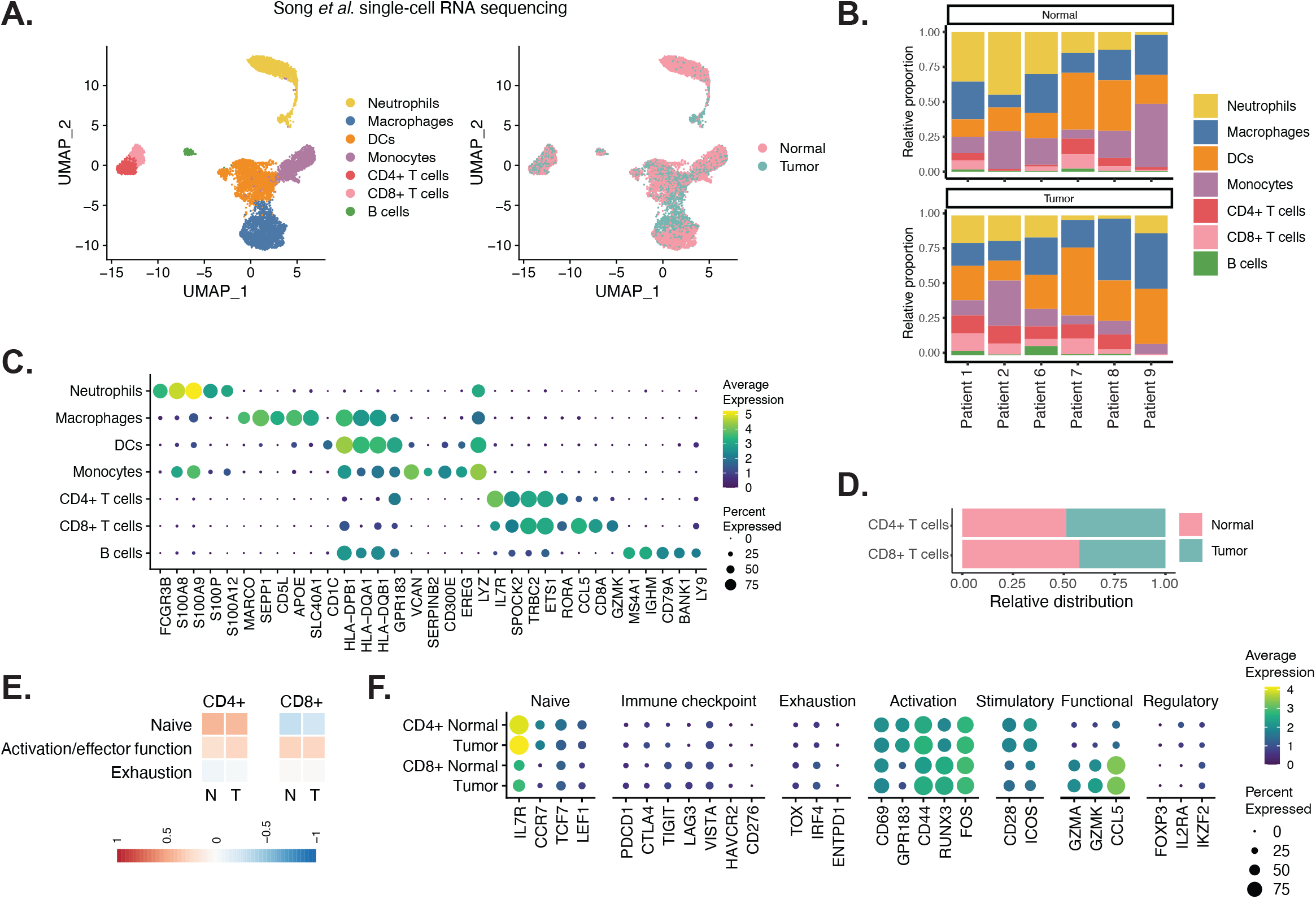
scRNA-seq shows presence of naive and early-activated T cells in chemotherapy-treated hepatoblastoma. **A.** UMAP of single cell transcriptome data of HB and distal normal tissues. Cells are colored by cell type (left) and tissue (right). **B.** Relative proportion of immune populations in normal and HB tissues per patient. Only patients from which more than 100 cells from each tissue were analyzed are displayed. **C.** Top 5 DEGs for each immune cluster **D.** Relative proportion of cells derived from normal or tumor tissue samples for each subcluster. **E.** Signature scores for T cell differentiation, split for normal (left) and tumor (right) samples. **F.** Expression of selected markers for different T cell states.

When comparing cell types from tumor and normal tissues, distinct transcriptional profiles were identified for macrophages, neutrophils, and B cells, but not for monocytes, DCs and T cells (Figure S8C,D). T cells were further subdivided into CD4^+^ and CD8^+^ subsets and were present in both normal and tumor tissue (Figure 5D). To further explore the phenotype of T cells, we analyzed T cell differentiation, based on gene signatures described by Chu et al.^25^ and expression of well-known markers for different T cell phenotypic states (Figure 5E,F, gene list in Table S3) ^26^. CD4^+^ T cells in normal and tumor tissues had a mixed phenotype characterized by high expression of naive (*IL7R*, *CCR7*, *TCF7, LEF1*) and tissue residency/activation (*CD69, GGPR183, CD44, RUNX3, FOS*) markers. CD8^+^ T cells displayed an early activated phenotype characterized by high expression of functional markers (*GZMA*, *GZMK* and *CCL5*). Of note, all T cells lacked expression of immune checkpoints (*PDCD1*, *CTLA4*, *TIGIT*, *LAG3, VISTA*), confirming IMC results. Moreover, all T cells lacked exhaustion markers *TOX, IRF4*, and *ENTPD1*. These results indicate the presence of naive and early activated T cells in both chemotherapy-treated HB and normal liver.

### HB macrophages express shared features with Kupffer cells

To further dissect the myeloid heterogeneity, a subset of cells was reclustered which identified two monocyte and two macrophage subpopulations (Figure 6A). The monocyte subpopulations were annotated as ‘CL monocytes’ and ‘intermediate (INT) and NC monocytes’, compared to a single monocyte cluster described by Song et al. ^12,27^ (Figure S8E, gene list in Table S4). Whereas no distinct transcriptional profiles were identified between tumor and normal tissue for both monocyte subsets, the transcriptional profiles of the two macrophage clusters were observed to be different (Figure S8D). One of the macrophage clusters was enriched in normal liver tissue and expressed KC markers (*MARCO*, *CD5L* and *TIMD4*), and was thus annotated ‘KC-like macrophages’ (Figure 6B,C). In normal tissue, the ‘KC-like macrophages’ highly expressed *MARCO*, demonstrating they are *bona fide* KCs (Figure 2E). The other macrophage cluster showed reduced expression or absence of KC genes, but expressed inflammatory markers such as *GPNMB*, *CCL18* and *SPP1*. The majority of these cells originated from tumor tissue and were annotated as ‘Inflammatory macrophages’. TAMs, from both clusters, displayed low *MARCO* expression, but expressed other KC markers (*CD5L*, *TIMD4*), with slightly higher expression in the ‘KC-like macrophages’ cluster (Figure 6D). In addition, TAMs showed increased levels of genes such as *APOE*, *CCL18* and *A2M*, which are abundantly expressed by ‘Inflammatory macrophages’. Furthermore, we scored the myeloid clusters for M1/M2 macrophage signatures and conserved gene signatures of tissue resident macrophages across tissues (gene lists in Table S5 and S6) ^21,28^. All normal and HB macrophage clusters showed an immunosuppressive M2 phenotype (Figure S8F). TAMs, found in both the inflammatory and KC-like clusters, expressed a high tissue residency signature compared to monocytes (Figure 6E). Taken together, these results indicate that tumor macrophages express inflammatory markers and, to a varying degree, KC and tissue residency markers.

**Figure 6:**
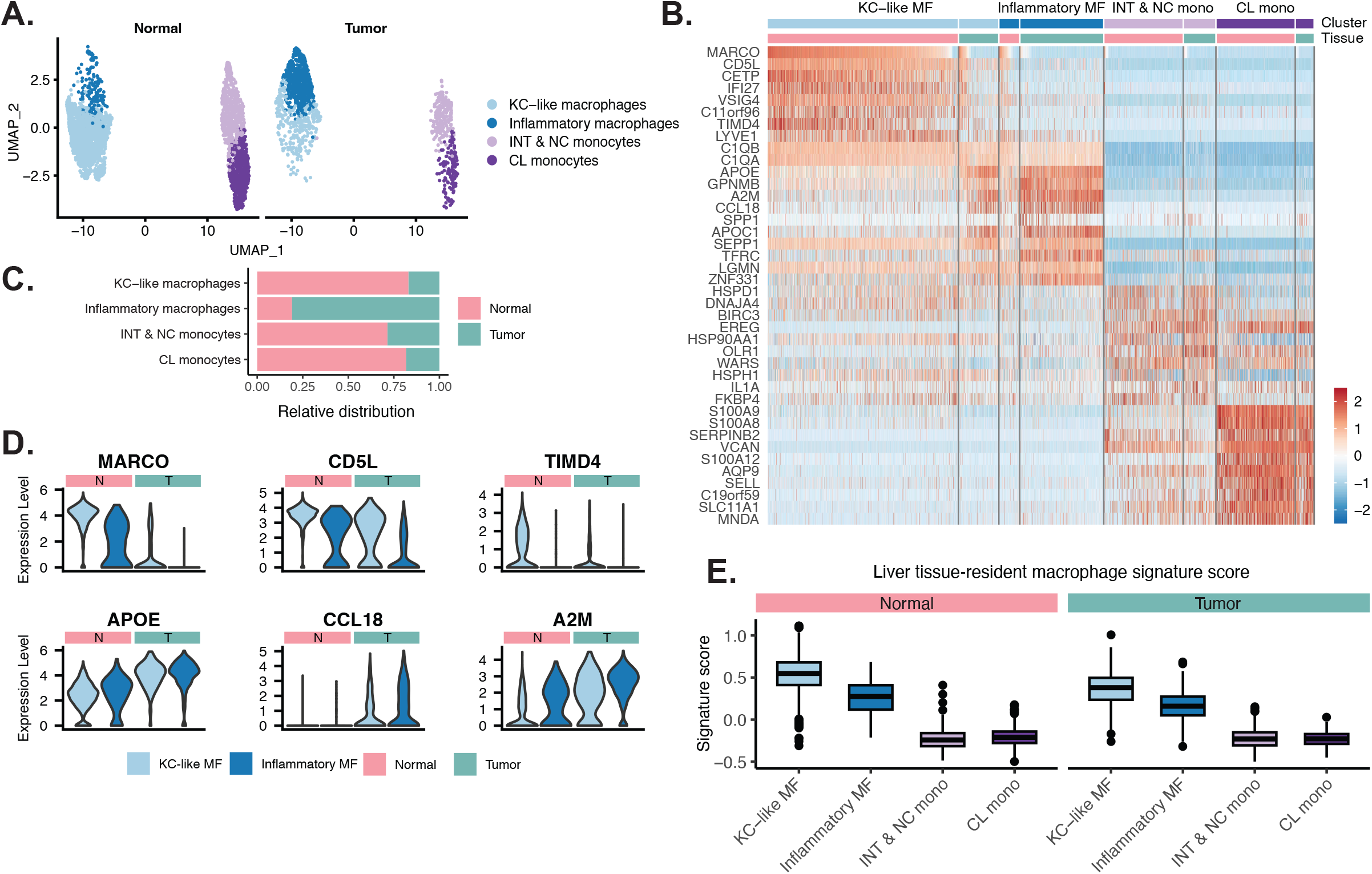
scRNA-seq of myeloid cells reveals expression of Kupffer cell and inflammatory markers in tumor-associated macrophages in hepatoblastoma. **A.** UMAP of macrophage and monocyte subclusters split by tumor and normal tissue. **B.** Relative proportion of cells derived from normal or tumor tissue samples for each subcluster. **C.** Top 10 DEGs for each subcluster split by tumor and normal tissue. **D.** Violin plot visualization of expression levels of selected cluster markers between KC-like macrophages from normal or tumor tissues. **E.** Signature scores for a liver tissue resident macrophage signature for macrophage and monocyte clusters, split for normal (left) and tumor (right) samples.

## Discussion

In this study, we characterized the immune landscape of HB using a synergistic approach that combines high-plex IMC and scRNA-seq to improve our understanding of the immunologic contribution to HB, thereby paving the way for future improvement in treatment options.

Currently, immune checkpoint inhibitors are being applied for treatment of advanced stage adult HCC ^10,29–31^. We demonstrate that tumor-associated T cells and T cells in distal normal liver of HB patients lack expression of immune checkpoint molecules, and do not display traits of exhaustion or dysfunction. This distinguishes HB tumors from adult HCC and emphasizes the challenges of using T cell targeting immune checkpoint blockade (ICB) in HB patients. Our findings are consistent with general observations that the expression of T cell immune checkpoints is rare in pediatric cancers, which are characterized by a low mutational burden and limited neoantigen expression ^32^.

Instead, we show that HB tumors are primarily infiltrated by myeloid immune cells which could be exploited for immunotherapeutic strategies. Although also observed by others ^12,33^, the current study provides the first detailed characterization of HB tissues in a spatial context, including investigation of immune checkpoint expression across different immune cell subsets. Macrophages are the primary effector cells in tissues performing antibody-dependent cellular phagocytosis (ADCP) via their Fcγ and Fcα receptors that bind IgG and IgA antibodies, respectively ^34,35^. A feasible therapeutic option might involve treating HB patients with therapeutic IgG and/or IgA antibodies directed against a tumor antigen which, in turn, activate macrophages in the TME to phagocytose the opsonized tumor cells. This poses two important challenges. First, a tumor-associated antigen is required. The fetal liver glycoprotein glypican-3 (GPC3) is one such antigen and is expressed in 95.5% of HB tumors ^36^. Anti-GPC3 IgG antibodies have been investigated in preclinical studies for adult HCC ^37–39^ and currently the first phase I trial is recruiting for relapsed or refractory HB (NCT04928677). Second, the presence of suppressive macrophages in the TME induces immune tolerance via inhibition of T-cell and NK-cell mediated responses, and differentiation of anti-tumor macrophages and T cells to suppressive M2 macrophages and Tregs, respectively ^40^. Moreover, pro-inflammatory macrophages can become suppressive upon ADCP via upregulation of immune checkpoints, thereby restraining the anti-tumor effects of macrophages ^41^. Together, it is likely that antibody therapy should be combined with a strategy to reactivate suppressive myeloid cells. We show that the suppressive ‘M2-like macrophages’ in HB express high levels of myeloid checkpoint molecules CD47, SIRPα, and VISTA. Therefore, myeloid checkpoint blockade with anti-CD47/SIRPα, or blockade of the recently identified myeloid checkpoint VISTA might be suitable treatment options for HB patients in combination with antibody therapy, which have been shown to restore phagocytic activity and shift macrophages to a pro-inflammatory M1 phenotype, respectively ^42,43^. Alternatively, new myeloid-targeting strategies including reprogramming of suppressive myeloid cells, and chimeric antigen receptor (CAR)-macrophages might be feasible strategies to treat HB in the future.

Several studies have demonstrated that human KCs can be identified on RNA level by expression of *MARCO*, *CD5L, VSIG4* and *TIMD4* ^21,44–47^. Our scRNA-seq analysis of chemotherapy-treated HB shows that TAMs have low *MARCO* expression and upregulated inflammatory markers, similar to transcriptional profiles of inflammatory monocyte-derived macrophages identified in several liver disease settings^30,46–50^. Surprisingly, we find that TAMs express KC markers (*CD5L, VSIG4, TIMD4*) and display high tissue residency scores in HB. It remains unclear whether these TAMs originate from KC, fetal macrophages, or monocytes, and what their role is in the TME. This also illustrates that additional heterogeneity within the myeloid TME needs to be considered when searching for novel immunotherapies targeting macrophages in HB.

Previous bulk RNA sequencing analyses demonstrated that chemotherapy could promote immune cell infiltration in HB ^13^. We reveal here that chemotherapy treatment induces specific influx of naive macrophages and active CD8^+^ T cells. Although immunomodulatory effects of chemotherapy have been described in literature ^51^, it is unclear which regimens are related to specific effects and this should therefore be explored in future studies. Interestingly, our transcriptomic analysis suggests that CD8^+^ T cells in chemotherapy-treated HB have an early activated phenotype. However, our neighborhood analysis did not show specific interactions between the infiltrated CD8^+^ T cells and naive macrophages with other cell types in chemotherapy-treated HB. It therefore remains unclear whether the infiltrated T cells and macrophages contribute to an anti-tumor response upon influx. Functional studies on these cells are needed to assess whether an actual anti-tumor response occurs.

This study is limited by the small sample size of eight tumor tissues and four normal liver tissues and was expanded with a scRNA-seq validation in a second dataset. Whereas the immune composition of normal tissue was homogeneous between patients, tumor tissue was highly heterogeneous. It is therefore likely that additional HB immune signatures exist that we were not able to capture in this study.

In summary, this study provides insights in the complex immune landscape of HB. Our results implicate unfeasibility of T cell-targeting with ICB in HB due to their low numbers, and absence of dysfunction and immune checkpoint expression. Instead, HB tumors contain a large suppressive myeloid compartment which poses opportunities for targeting. Specifically, therapeutic antibodies targeting the tumor antigen GPC3 to induce ADCP by macrophages, in combination with myeloid immune checkpoint blockade (CD47/SIRPa, or VISTA) could be a feasible strategy to explore in future studies.

## Abbreviations

ADCP: Antibody-dependent cellular phagocytosis
APC: Antigen presenting cells
BSA: Bovine serum albumin
CAR: Chimeric antigen receptor
CL: Classical
FFPE: Formalin-fixed paraffin-embedded
GPC3: Glycoprotein glypical-3
HB: Hepatoblastoma
HCC: Hepatocellular carcinoma
ICB: Immune checkpoint blockade
IF: Immunofluorescence
IHC: Immunohistochemistry
IMC: Imaging mass cytometry
INT: Intermediate
KC: Kupffer cell
NC: Non-classical
P-ERK: Phosphorylated ERK
PCA: Principal component analysis
PMC: Princess Máxima Center
RNA-seq: RNA sequencing
ROI: Region of interest
RT: Room temperature
scRNA-seq: Single-cell RNA-sequencing
TAM: Tumor-associated macrophages
TMA: Tissue microarray
TME: Tumor microenvironment
UMAP: Uniform manifold approximation and projection
UMCG: University Medical Center Groningen
UMCU: University Medical Center Utrecht

## Acknowledgements

We would like to thank the physicians and research nurses who are part of the Máxima Comprehensive Childhood Cancer Center (M4C) liver tumor group, especially Kathelijne Kraal, Martine van Grotel, Kees van de Ven, Lideke van der Steeg, Liset Lansaat (Princess Máxima Center) and Frank Bodowes (UMCG). We thank Livio Kleij (Center for Molecular Medicine, University Medical Center Utrecht; UMCU), and Domenico Castigliego (Department of Pathology, UMCU) for technical support. Furthermore, we thank Mojtaba Amini, Pascalle Hemelop, Dorotea Neuberg, and Pilar van Wijngaarden (Center for Molecular Medicine, UMCU) for practical support with antibody labeling and validation. Additionally, we thank UCyTOF (UMCU) for support. Finally, we thank Julia Drylewics and Laura Guerrero Simon (Computational Immunology Core, UMCU) for statistical advice and division LAB ICT team for infrastructural support.

