## Supplemental material for "Hepatoblastoma exhibits a predominantly myeloid immune landscape and reveals opportunities for macrophage targeted immunotherapy"

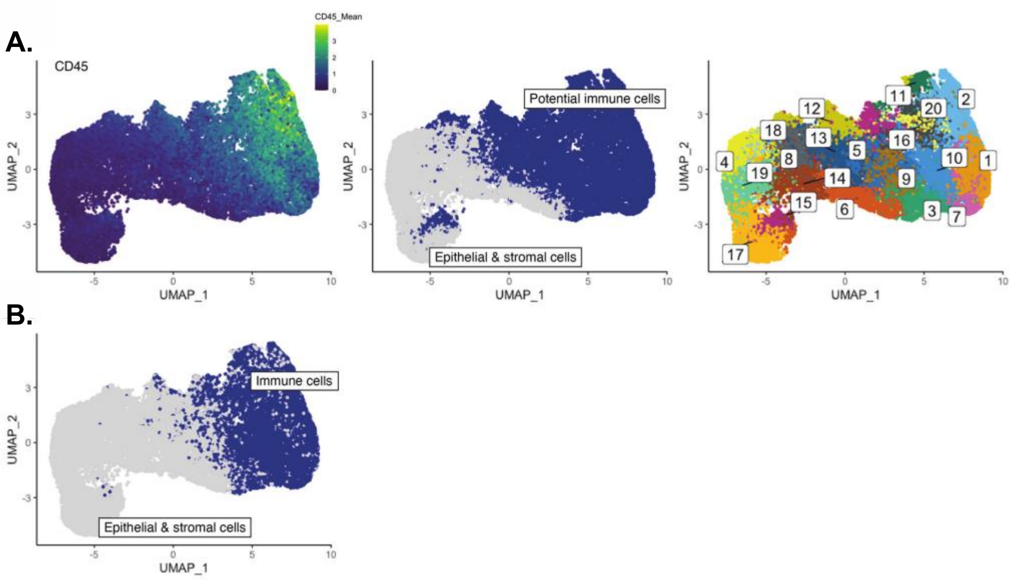


**Figure S1: UMAPs of CD45⁺ cell clustering**

**A.** Uniform Manifold Approximation and Projection (UMAP) of single cells of HB tissues and normal tissues together, with projection of CD45 expression (left), annotation of potential immune cells (center), and identified clusters (right). **B.** Annotated UMAP with immune and epithelial & stromal clusters after additional clustering as indicated in Figure 2 and 3.

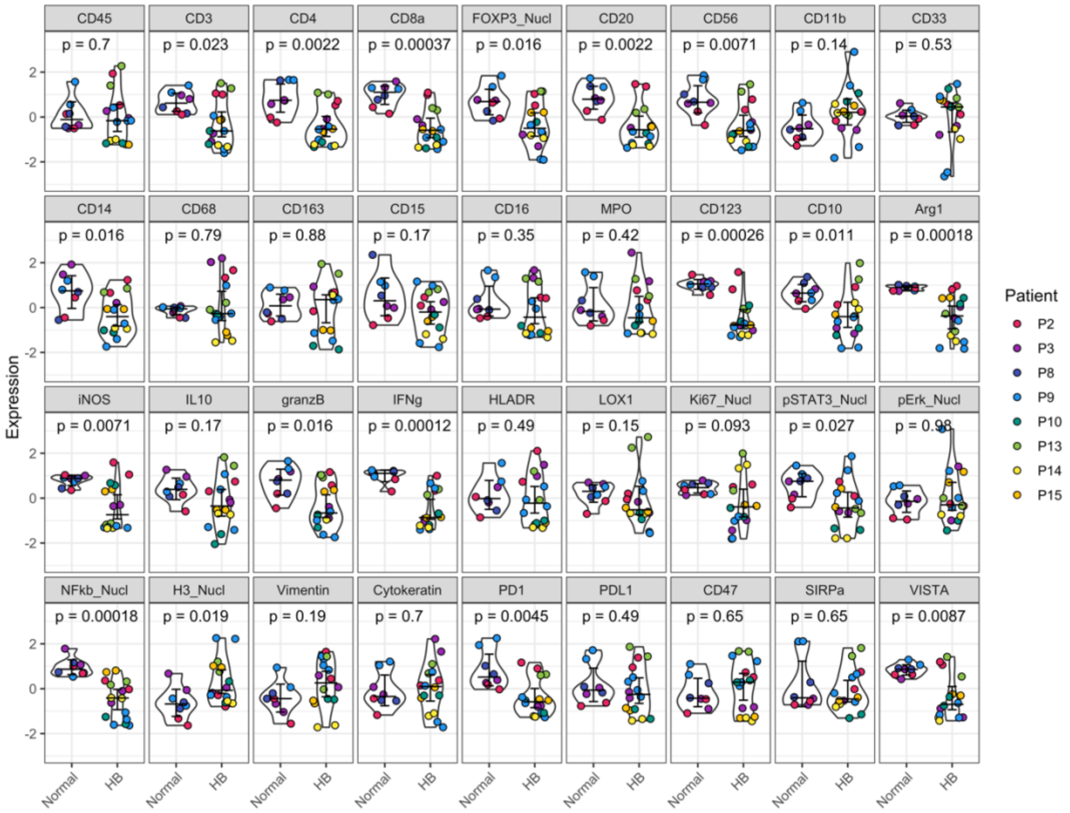


**Figure S2: Immune checkpoint expression in hepatoblastoma and adjacent normal tissue**

Violin plots with median intensity of immune markers per patient, compared between HB tissues and adjacent normal liver. Patient ID is indicated in colors. Nonparametric T tests were used to calculate the statistical difference between HB and normal tissues. Bars indicate the median with 95% confidence interval.

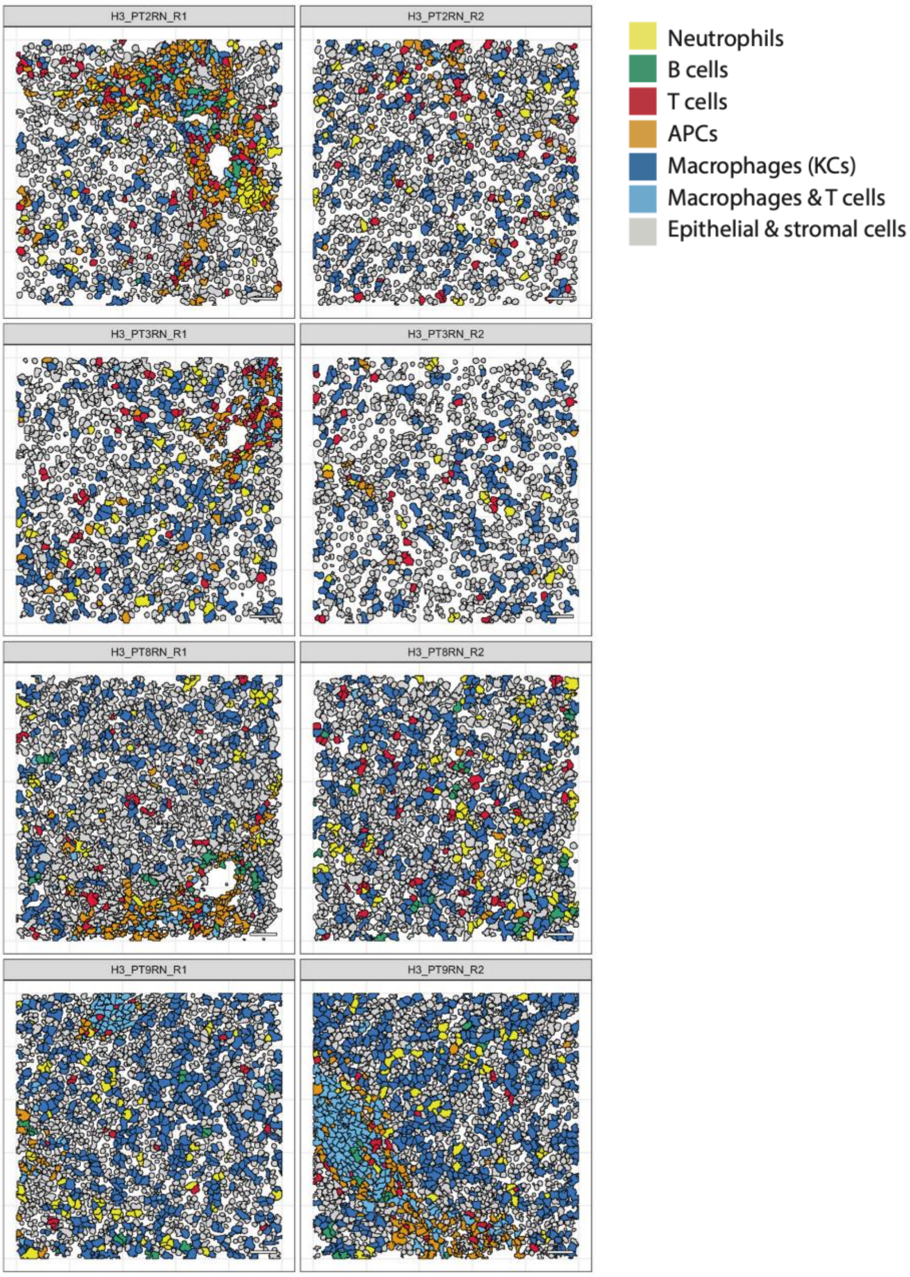


**Figure S3: Single-cell segmented cells of normal liver tissues**

Single-cell segmented cells of all normal liver tissues included in this study. Colors indicate the annotated cell types. Scale bar is 50 μm.

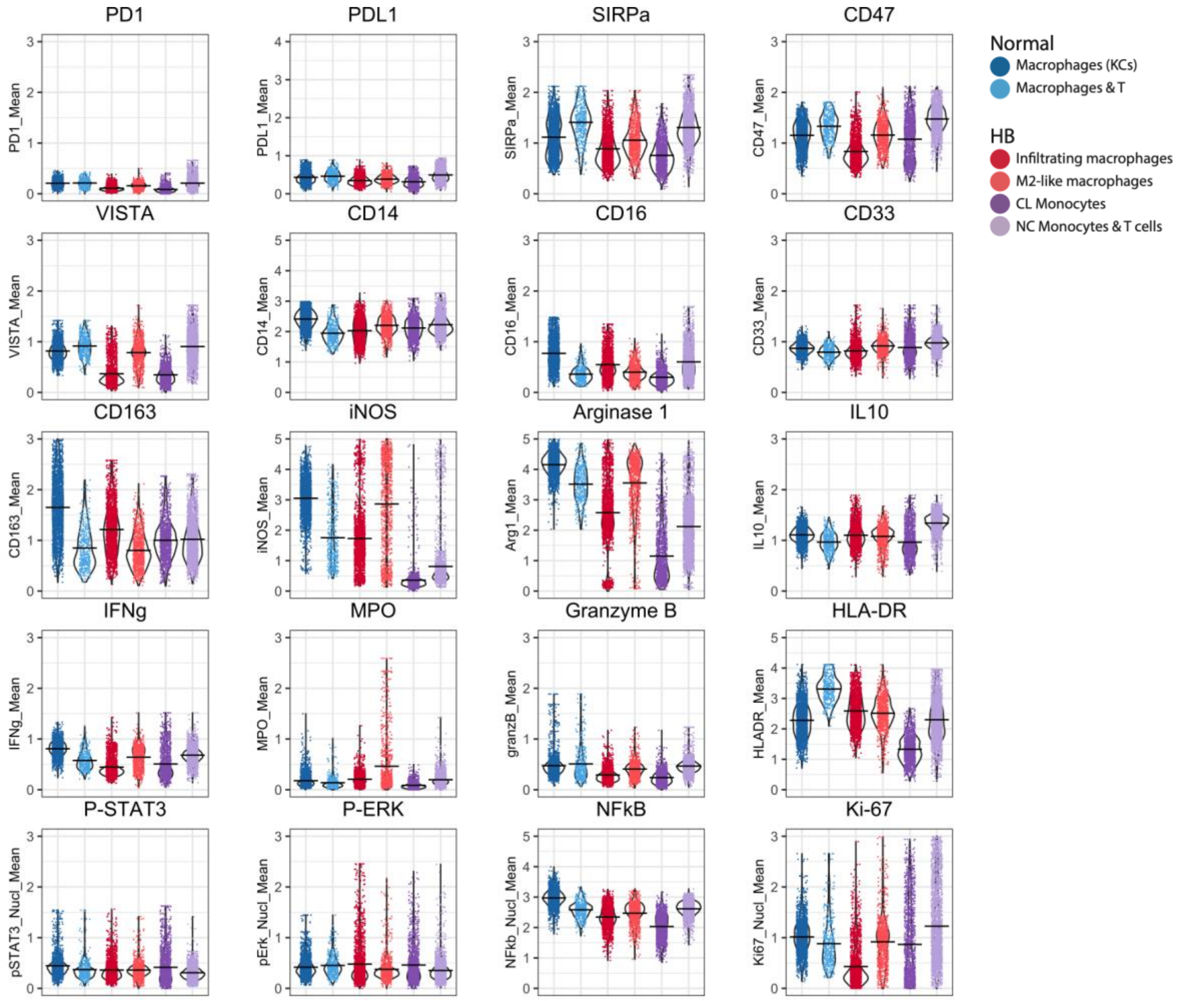


**Figure S4: Phenotype of myeloid cell clusters**

Scatter plot of immune checkpoint expression, functional markers and signaling markers across myeloid clusters in HB and normal samples. The bars indicate the median with 95% confidence interval.

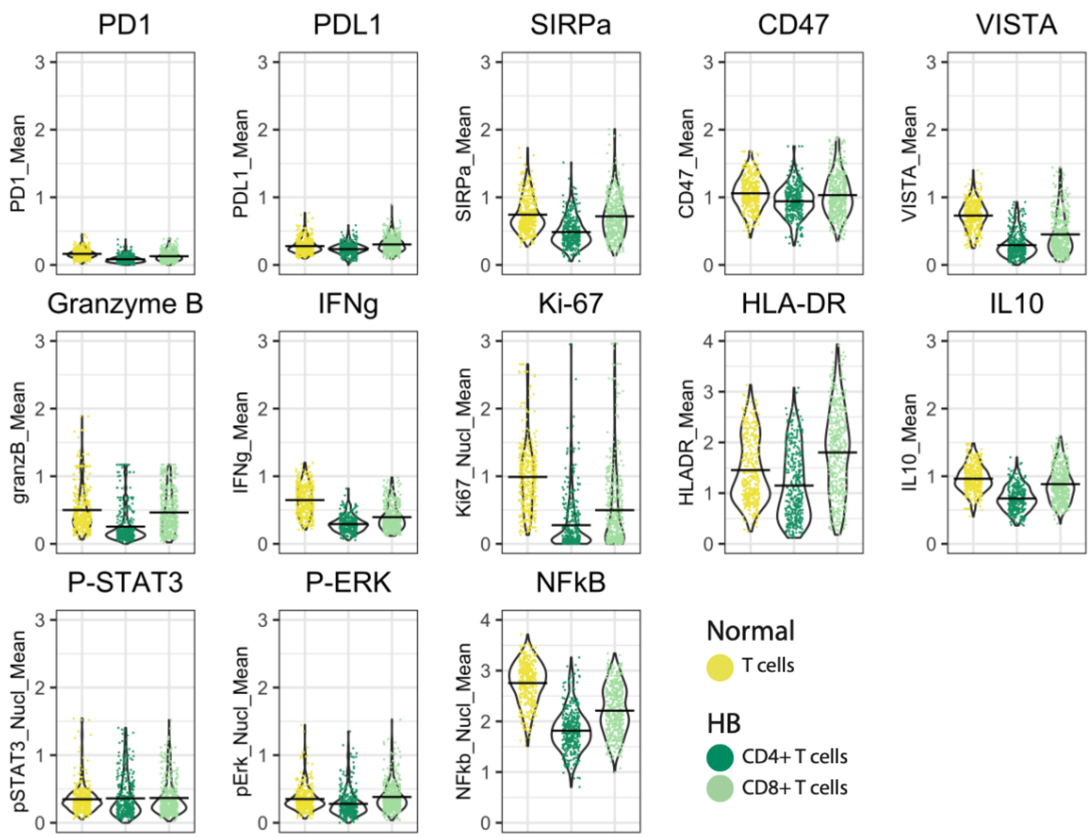


**Figure S5: Phenotype of T cell clusters**

Scatter plot of immune checkpoint expression, functional markers and signaling markers across T cell clusters in HB and normal samples. The bars indicate the median with 95% confidence interval.

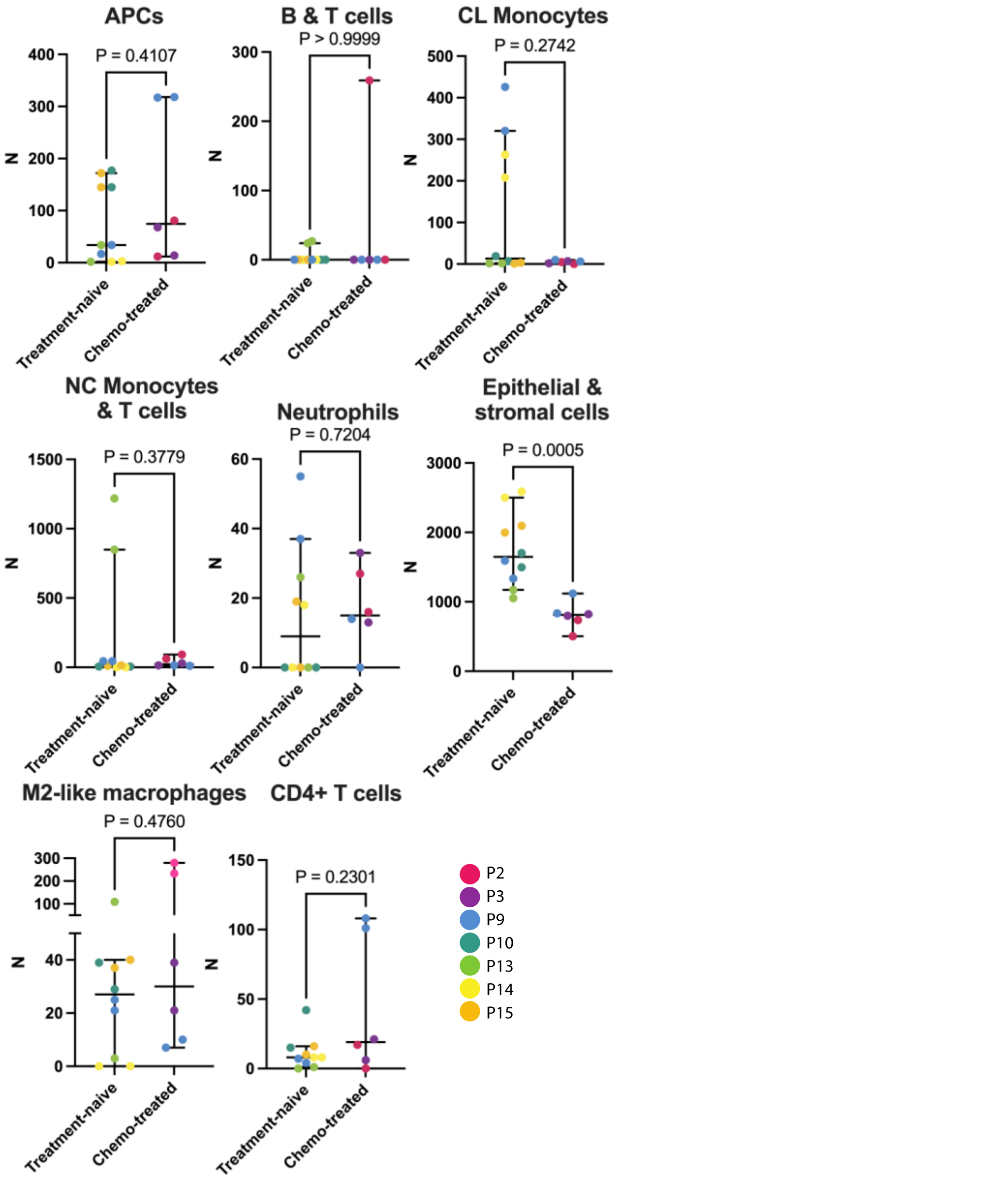


**Figure S6: Immune cluster abundancy in hepatoblastoma tissues.**

Scatter plots of the number of cells per annotated cell cluster, stratified by treatment status. Each dot is an ROI, color-coded for patient ID. The bars indicate the median with 95% confidence interval. Nonparametric T tests were used to calculate the statistical difference between abundance of cell clusters in treatment-naive and chemotherapy-treated HB samples.

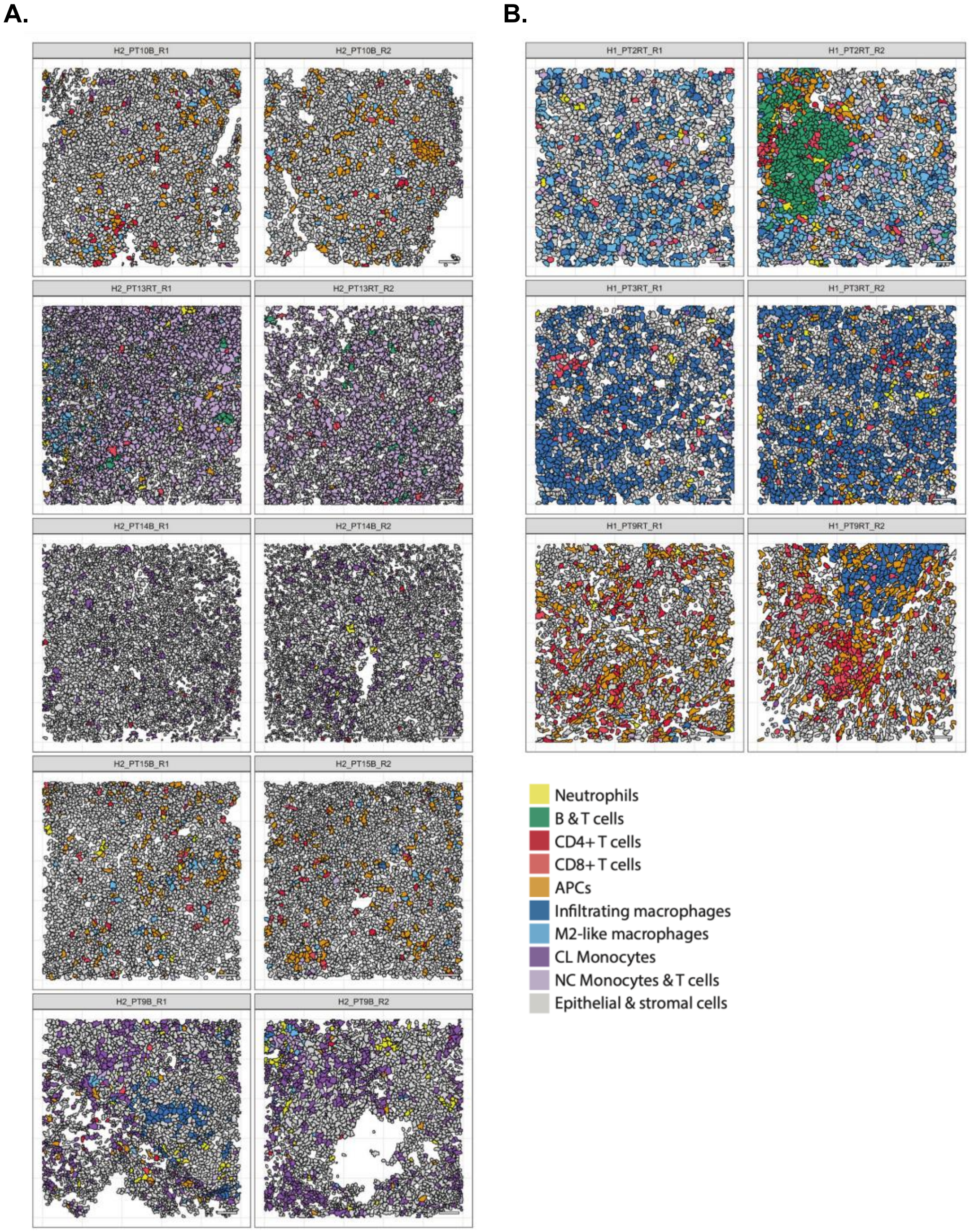


**Figure S7: Single-cell segmented cells of hepatoblastoma tissues**

Single-cell segmented cells of treatment-naive (**A.**) and chemotherapy-treated (**B.**) HB tissues included in this study. Colors indicate the annotated cell types. Scale bar is 50 μm.

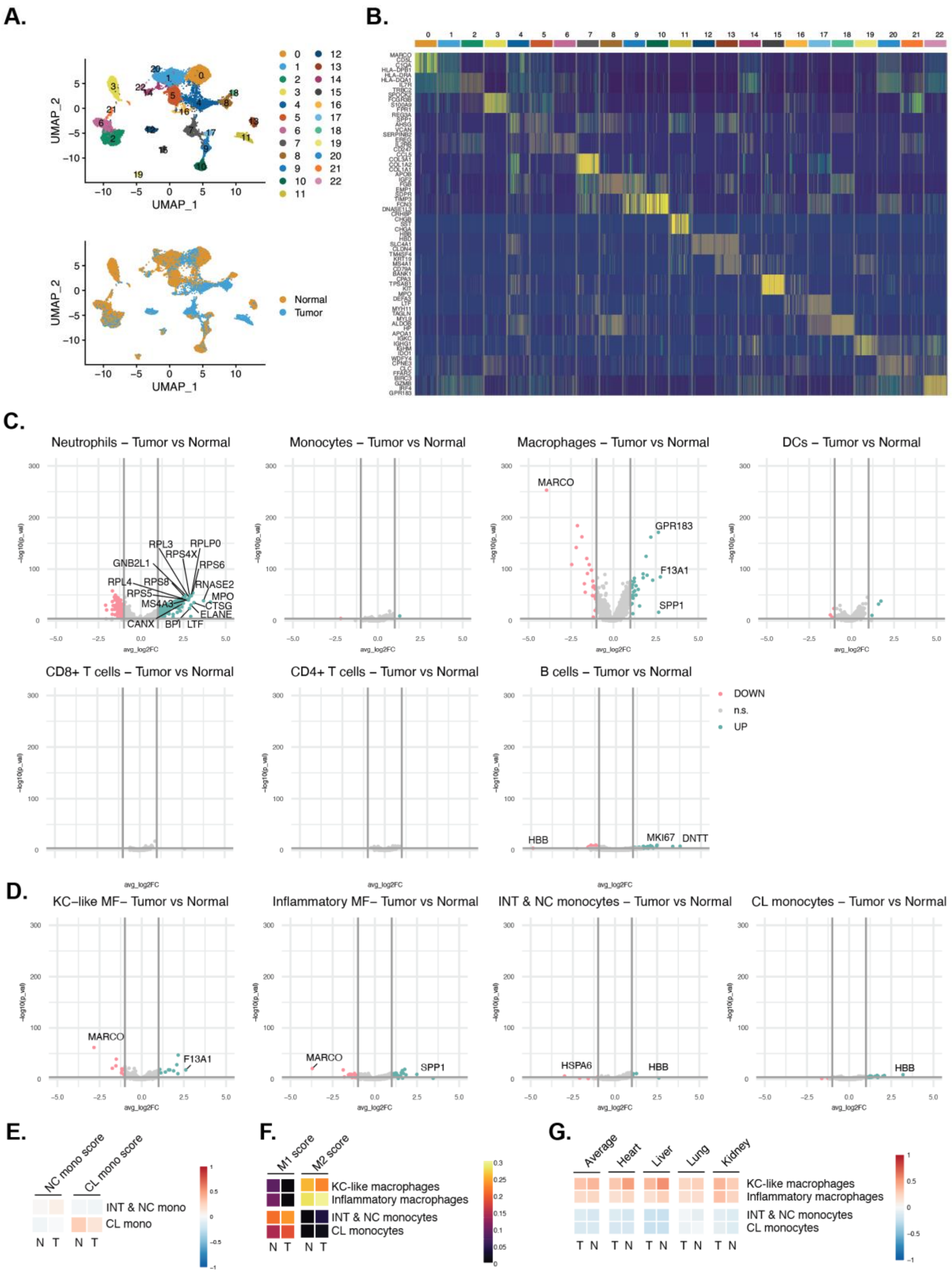


**Figure S8: Extended single-cell RNA sequencing data**

**A.** Clustering of cells from chemotherapy-treated HB and adjacent normal tissues from 9 patients into 22 clusters. **B.** Top 3 differentially expressed genes per cluster. **C.** Differentially expressed genes between tumor and normal cells within the same immune cluster. **D.** Differentially expressed genes between tumor and normal cells within the same myeloid subcluster. **E.** Signature scores for monocyte subset specific signatures per cluster split by tumor and normal cells. **F.** Signature scores for M1 and M2 cluster split by tumor and normal cells. **G.** Signature scores for tissue-resident macrophages of multiple organs and the average score for all organs per cluster split by tumor and normal cells.

**Table S1: Antibody list for IMC**

| Mass | Metal | Marker | Clone | Company | Dilution |
|---|---|---|---|---|---|
| **140** | Ce | CD10 | E5P7S | CST | 1:50 |
| **141** | Pr | CD56 | NCAM1/784 | Abcam | 1:100 |
| **142** | Nd | Cytokeratin | C11 | CST | 1:100 |
| **143** | Nd | MPO | E1E7I | CST | 1:200 |
| **144** | Nd | CD16 | EPR22409-124 | Abcam | 1:50 |
| **145** | Nd | CD33 | Polyclonal | Fluidigm | 1:75 |
| **146** | Nd | CD15 | W6D3 | Biolegend | 1:200 |
| **147** | Sm | CD163 | D6U1J | CST | 1:200 |
| **148** | Nd | PD-L1 | E1L3N | CST | 1:50 |
| **149** | Sm | Vimentin | D21H3 | CST | 1:100 |
| **150** | Nd | CD47 | EPR21794 | Abcam | 1:100 |
| **151** | Eu | IFNg | D3H2 | CST | 1:100 |
| **152** | Sm | iNOS | SP126 | Abcam | 1:50 |
| **153** | Eu | NFkB | L8F6 | CST | 1:200 |
| **154** | Sm | CD68 | KP1 | Biolegend | 1:200 |
| **155** | Gd | FOXP3 | 236A/E9 | Abcam | 1:50 |
| **156** | Gd | CD4 | EPR6855 | Abcam | 1:50 |
| **158** | Gd | pSTAT3 | D3A7 | CST | 1:200 |
| **159** | Tb | CD20 | H1 | BD biosciences | 1:200 |
| **160** | Gd | IL-10 | Polyclonal | R&D | 1:200 |
| **161** | Dy | Secondary rabbit | Polyclonal | Invitrogen | 1:100 |
| **162** | Dy | LOX-1 | Polyclonal | Invitrogen | 1:100 |
| **163** | Dy | CD123 | IL3RA/1531 | LsBio | 1:100 |
| **164** | Dy | CD45 | D9M8I | CST | 1:200 |
| **165** | Ho | PD-1 | EPR4877(2) | Abcam | 1:50 |
| **166** | Er | VISTA | D1L2G | CST | 1:100 |
| **167** | Er | Arginase 1 | SI6arg | eBiosciences | 1:400 |
| **168** | Er | Ki-67 | B56 | BD Biosciences | 1:200 |
| **169** | Tb | Granzyme B | D6E9W | CST | 1:200 |
| **170** | Er | CD3 | Polyclonal | DAKO | 1:100 |
| **171** | Yb | pERK [T202/Y204] | D13.14.4E | CST | 1:100 |
| **172** | Yb | CD8a | C8/144B | ThermoFisher | 1:200 |
| **173** | Yb | HLA-DR | TAL 1B5 | Abcam | 1:800 |
| **174** | Yb | CD14 | EPR3653 | Abcam | 1:200 |
| **175** | Lu | CD11b | D6X1N | CST | 1:200 |
| **176** | Yb | Histon H3 | D1H2 | CST | 1:800 |
| | | SIRPa | D6I3M | CST | 1:100 |

**Table S2: Antibody list for IF Kupffer cell staining**

| Antibody | Supplier | Cat.No. | Dilution |
|---|---|---|---|
| MARCO | Sigma Aldrich | HPA063793 | 1:100 |
| CD68 | Leica Biosystems | NCL-L-CD68 | 1:100 |
| Anti-rabbit 555 | ThermoFisher | A-31572 | 1:1000 |
| Anti-mouse 647 | ThermoFisher | A-31571 | 1:1000 |

**Table S3: Human genes for T-cell differentiation scores**

(Chu et al. Pan-cancer T cell atlas links a cellular stress response state to immunotherapy resistance. Nat Med. 2023)

| CD4+ naïve | CD4+ activation/effector | CD4+ exhaustion | CD8+ naive | CD8+ activation/effector | CD8+ exhaustion |
|---|---|---|---|---|---|
| IL7R | FAS | PDCD1 | IL7R | FAS | PDCD1 |
| CCR7 | CD44 | LAYN | CCR7 | FASLG | LAYN |
| SELL | CD69 | HAVCR2 | SELL | CD44 | HAVCR2 |
| FOXP1 | CD38 | LAG3 | FOXO1 | CD69 | LAG3 |
| KLF2 | NKG7 | CTLA4 | KLF2 | CD38 | CD244 |
| KLF3 | KLRB1 | TIGIT | KLF3 | NKG7 | CTLA4 |
| LEF1 | KLRD1 | TOX | LEF1 | KLRB1 | LILRB1 |
| TCF7 | KLRG1 | VSIR | TCF7 | KLRD1 | TIGIT |
| ACTN1 | CX3CR1 | BTLA | ACTN1 | KLRF1 | TOX |
| BTG1 | CD300A | ENTPD1 | FOXP1 | KLRG1 | VSIR |
| BTG2 | FGFBP2 | | | KLRK1 | BTLA |
| TOB1 | ID2 | | | FCGR3A | ENTPD1 |
| | ID3 | | | CX3CR1 | CD160 |
| | PRDM1 | | | CD300A | LAIR1 |
| | RUNX3 | | | FGFBP2 | |
| | TBX21 | | | ID2 | |
| | ZEB2 | | | ID3 | |
| | BATF | | | PRDM1 | |
| | NR4A1 | | | RUNX3 | |
| | NR4A2 | | | TBX21 | |
| | HOPX | | | ZEB2 | |
| | FOS | | | BATF | |
| | FOSB | | | IRF4 | |
| | FOSL2 | | | NR4A1 | |
| | JUN | | | NR4A2 | |
| | JUNB | | | NR4A3 | |
| | JUND | | | PBX3 | |
| | STAT1 | | | ZNF683 | |
| | STAT3 | | | HOPX | |
| | EOMES | | | FOS | |
| | AHR | | | FOSB | |
| | | | | JUN | |
| | | | | JUNB | |
| | | | | JUND | |
| | | | | STAT1 | |
| | | | | STAT2 | |
| | | | | STAT5A | |
| | | | | STAT6 | |
| | | | | STAT4 | |
| | | | | EOMES | |

**Table S4: Human genes for monocyte scores**

(Vallania et al. Multicohort Analysis Identifies Monocyte Gene Signatures to Accurately Monitor Subset-Specific Changes in Human Diseases. Front Immunol. 2021)

| CL monoyctes | NC monocytes |
| --- | --- |
| PGD | TCF7L2 |
| FCGR2A | IER2 |
| MS4A6A | CDKN1C |
| CSF3R | SIGLEC10 |
| IL17RA | NFKBIZ |
| DPYD | NAP1L1 |
| ANPEP | FCGR3B |
| CD36 | TOB1 |
| ATP6V0E1 | RP4-781L3.1 |
| LMAN2 | CSTA |

**Table S5: Human genes for M1 and M2 scores**

(Cheng et al. A pan-cancer single-cell transcriptional atlas of tumor infiltrating myeloid cells. Cell. 2021).

| M1 | M2 |
|---|---|
| IL23 | IL4R |
| TNF | CCL4 |
| CXCL9 | CCL13 |
| CXCL10 | CCL20 |
| CXCL11 | CCL17 |
| CD86 | CCL18 |
| IL1A | CCL22 |
| IL1B | CCL24 |
| IL6 | LYVE1 |
| CCL5 | VEGFA |
| IRF5 | VEGFB |
| IRF1 | VEGFC |
| CD40 | VEGFD |
| IDO1 | EGF |
| KYNU | CTSA |
| CCR7 | CTSB |
| | CTSC |
| | CTSD |
| | TGFB1 |
| | TGFB2 |
| | TGFB3 |
| | MMP14 |
| | MMP19 |
| | MMP9 |
| | CLEC7A |
| | WNT7B |
| | FASL |
| | TNFSF12 |
| | TNFSF8 |
| | CD276 |
| | VTCN1 |
| | MSR1 |
| | FN1 |
| | IRF4 |

**Table S6: Human orthologs of mouse gene for macrophage tissue residency scores**

(Dick et al. Three tissue resident macrophage subsets coexist across organs with conserved origins and life cycles. Sci Immunol. 2022)

| Heart TLF | Heart CCR | Liver TLF | Liver CCR | Lung TLF | Lung CCR | Kidney TLF | Kidney CCR |
|---|---|---|---|---|---|---|---|
| LYVE1 | CD9 | CD5L | PLAC8 | CLEC4M | MMP12 | CXCL13 | CCR2 |
| FOLR2 | CD74 | APOC1 | TMSB10 | F13A1 | LGALS3 | FOLR3 | MMP12 |
| FOLR3 | HLA-DQB1 | VSIG4 | CXCL9 | CCL24 | S100A6 | FOLR2 | NAPSA |
| F13A1 | HLA-DQB2 | CLEC4F | S100A6 | CLEC10A | CCR2 | PLA2G2D | TMSB10 |
| CCL23 | CCR2 | HMOX1 | BCL2A1 | FOLR3 | C15orf48 | CCL2 | CLEC4C |
| CCL15 | HLA-DMB | FABP7 | S100A4 | FOLR2 | TPPP3 | CLEC4M | S100A6 |
| RETNLB | CD52 | SLC40A1 | S100A11 | LYVE1 | CD9 | F13A1 | CLEC12A |
| NINJ1 | CD72 | FOLR3 | GBP2 | C4B | LSP1 | NINJ1 | ITGAX |
| VSIG4 | HLA-DMA | FOLR2 | LSP1 | C4A | HLA-C | MRC1 | CCL23 |
| CCL24 | GNGT2 | CD163 | CRIP1 | CD209 | HLA-E | STAB1 | CCL15 |
| FXYD2 | HEXB | SDC3 | CHIA | CD163 | HLA-G | IGF1 | CD9 |
| FCGRT | TMSB10 | C6 | NAPSA | EDNRB | HLA-F | APOE | CYP4F3 |
| SELENBP1 | SCIMP | CFP | VIM | GAS6 | HLA-A | GAS6 | FCGR3B |
| TIMD4 | MPEG1 | CDH5 | CCL23 | NINJ1 | HLA-B | IGFBP4 | FCGR3A |
| KLF2 | AXL | TIMD4 | CCL15 | RCN3 | CRIP1 | CFP | PSAP |
| CD163 | CXCL14 | MRC1 | CCR2 | FCGRT | S100A4 | LY6E | SIRPG |
| PLTP | PLBD1 | CCL24 | C3 | CCL23 | CTSS | CTSD | SIRPB1 |
| RCN3 | TGFBR1 | C4B | CXCL10 | CCL15 | CD74 | MAF | SIRPA |
| CFP | FCGR3B | C4A | MS4A4A | FXYD2 | BCL2A1 | | TGFBR1 |
| GAS6 | FCGR3A | NR1H3 | MS4A4E | CD36 | TMSB10 | | LPL |
| PEPD | CXCL16 | DMPK | S100A10 | MAF | CAPG | | FYB1 |
| | CKB | PAQR9 | TSPO | IFITM3 | HLA-DMB | | |
| | GLIPR1 | IL18BP | | IFITM2 | HLA-DQB1 | | |
| | MMP13 | CD164 | | IFITM1 | HLA-DQB2 | | |
| | | C6orf62 | | | NAPSA | | |
| | | ACTN1 | | | F11R | | |
| | | | | | PSAP | | |
